# Muscleblind-like proteins dimerize by forming disulfide bonds to regulate alternative splicing and pathogenic RNA foci formation

**DOI:** 10.64898/2026.03.24.714019

**Authors:** Luke A. Knudson, Adam Kosti, Kathryn R. Moss, Liang Shi, GiaLinh N. Nguyen, Aleksandra Janusz-Kaminska, Eric X. Zhou, Ryan P. Hildebrandt, Eric T. Wang, Gary J. Bassell

## Abstract

Muscleblind-like (MBNL) RNA-binding proteins (RBPs) possess modular domains that mediate regulation of alternative splicing and RNA localization. Myotonic Dystrophy Type 1 is a CTG repeat expansion disorder where MBNL is sequestered into intranuclear RNA foci, impairing its function. Previous studies found that MBNL self-associates through its exon 7, but the nature of this interaction is not well understood. We identified a cysteine in MBNL1 exon 7 that enables dimerization through formation of an intermolecular disulfide bond. We likewise demonstrate that MBNL2 dimerizes by forming disulfide bonds between multiple cysteines in its carboxy-terminus. Nucleocytoplasmic fractionation revealed a greater proportion of MBNL1 dimer in the nucleus, suggesting a nuclear function for the MBNL1 dimer. We investigated a connection between MBNL1 dimerization and MBNL1-mediated regulation of alternative splicing. To accomplish this, we mutated the MBNL1 cysteine in question to alanine (C325A) and performed RNAseq. We uncovered novel splicing events sensitive to MBNL1 dimerization. We also found that MBNL1 C325A, when co-expressed with expanded CTG repeats, produces smaller, more numerous foci, suggesting a role for the MBNL1 dimer in maintaining foci integrity. These results provide insight into biological and pathological mechanisms of MBNL1 dimerization and suggest other RBPs might similarly dimerize to regulate function.

**GRAPHICAL ABSTRACT:** 

## INTRODUCTION

RNA-binding proteins (RBPs) frequently cooperate to bind target transcripts containing multiple cognate motifs, allowing for proper and efficient regulation of alternative splicing (1-4). This “multivalency” often arises through interactions between RBPs involving intrinsically disordered domains (IDRs) (5). IDRs have been shown to mediate both homomeric and heteromeric interactions in a range of splicing regulators, including SR and SR-related proteins (5), RBFOX2 (6), hnRNPH1 (7), RBPMS (8), hnRNPA and hnRNPD (9). While RBP-RBP interactions can play a physiological role in processes such as splicing, they can also contribute to disease pathology. Aberrant expansions of repetitive sequences in the human genome have been linked to multiple diseases. In many of these diseases, the RNA transcribed from expanded repeats accumulates within cells, forming RNA-protein aggregates, called RNA foci. Besides RNA-RNA and RNA-RBP interactions, RBP-RBP interactions have been proposed as a factor that stabilizes RNA foci (10). Whether in a physiologic or pathologic context, the relatively weak non-covalent interactions produced by IDRs are not the only forces capable of bringing RBPs together (11). Intermolecular disulfide bonds have been found to form between cysteine residues located in several RBPs, including TDP-43, FMRP, and PTB (12-14). However, compared to non-covalent interactions involving IDRs, disulfide bond formation between RBPs has been understudied, especially as it relates to the effect it has on RBP function.

The Muscleblind-like (MBNL) family of RBPs are known for their capacity to regulate alternative splicing (15-17), alternative polyadenylation (18), mRNA localization (17,19,20), translation (17), and mRNA stability (21). MBNL itself undergoes extensive alternative splicing, producing isoforms found in both the nucleus and cytoplasm, depending on the presence of a full or partial bipartite nuclear localization signal (NLS), respectively. In the nucleus, MBNL proteins regulate alternative splicing using two pairs of zinc fingers that bind a consensus motif (22-24), YGCY (15-17), in target pre-mRNA. In the cytoplasm, we found that MBNL binding to 3’UTRs localizes target mRNAs by interacting with specific kinesin motor proteins (20). Besides two pairs of zinc fingers, MBNL proteins also possess an unstructured C-terminal domain, which facilitates anchoring of MBNL to membranes (20). In addition to membrane anchoring, multiple studies have demonstrated that the C-terminus (25), specifically exon 7, enables MBNL1 to interact with itself (26,27), forming an MBNL1 dimer. Exon 7 has also emerged as a modulator of several MBNL1-dependent alternative splicing events (28,29), but this has not been linked to exon 7-dependent MBNL self-association. Furthermore, the nature of the bond holding two MBNL1 molecules together and the potential impact this has on MBNL1-mediated alternative splicing regulation are both unknown.

MBNL proteins play a prominent role in the pathogenesis of the repeat expansion disorder Myotonic Dystrophy Type 1 (DM1) (30,31). In DM1, expanded CTG repeats in the 3’UTR of the gene *DMPK* are transcribed into toxic RNA that sequesters MBNL proteins in the nucleus, forming CUG RNA foci (32-34). Sequestered MBNL is nonfunctional, resulting in an absence of MBNL-dependent splicing regulation in individuals with DM1 (35). Despite the importance of MBNL1 in the pathophysiology of DM1, the impact of MBNL1 dimerization on DM1 remains unexplored.

In this study, we utilized immunoprecipitation (IP) to confirm the existence of MBNL1 dimers and determined that they are sensitive to reducing agents. IP also revealed the existence of reducing agent-sensitive MBNL2 dimers. Taking these observations into consideration, we hypothesized that at least one disulfide bond holds both MBNL1 and MBNL2 homodimers together. To test this, we mutated specific cysteine residues in MBNL1 and MBNL2, performed IPs, and assessed the effect the mutations had on the ability of MBNL to dimerize. In the process, we found that IP was unnecessary to visualize MBNL1 dimers extracted from the nucleus. The increased prevalence suggested a potential nuclear function for the MBNL dimer, such as in the regulation of alternative splicing. We used *Mbnl1*^*-/-*^; *Mbnl2*^*-/-*^ mouse embryonic fibroblasts (DKO MEFs) (18,31), a MBNL1 cysteine mutant (C325A), and RNA-seq to investigate the effect of MBNL1 dimerization on MBNL1-dependent alternative splicing regulation. Finally, to uncover a possible role for MBNL1 dimerization in DM1 pathogenesis, we imaged CUG foci in DKO MEFs co-transfected with CTG repeats and either wildtype or C325A GFP-MBNL1. Findings from this study suggest that MBNL1 dimerization plays a physiological role in RNA biology and has pathological consequences for DM1 pathogenesis.

## MATERIAL AND METHODS

### Plasmids and Cloning

Human *MBNL1* and *MBNL2* isoforms were amplified from cDNA constructs obtained from the DNASU Plasmid Repository (Tempe, AZ) or GenScript (Piscataway, NJ) and subcloned downstream of EGFP under control of a CMV promoter (pEGFP-C1; Takara). MBNL1 cysteine mutants (C325A [41] and C343A [43]) were generated by Genewiz and the MBNL2 cysteine mutants (C324A [I1], C332 [I3], C343A [I3], C345A [I3], and C361A [I3]) were generated by GenScript. Plasmids carrying exons 11–15 of the *DMPK* gene expressing 0 (DMPKS) or 480 CTG repeats under the control of a CMV promoter were a gift from Dr. Thomas Cooper (Baylor College of Medicine). The *TNNT2* minigene (RTB300) was also a gift from Dr. Thomas Cooper (Addgene plasmid #80410; http://n2t.net/addgene:80410; RRID:Addgene_80410) (36).

### Cell Lines and Transfection

Neuro2a cells (ATCC, CCL-131), SH-SY5Y cells (ATCC, CRL-2266), and MBNL1/2 double-knockout mouse embryonic fibroblasts (DKO MEFs; gift from Dr. Maurice Swanson) were cultured in DMEM, high glucose, pyruvate supplemented with 100 U/mL penicillin, 100 mg/mL streptomycin, 10mM HEPES, and 10% fetal bovine serum (FBS). Wildtype and DM1 human fibroblasts (gift from Dr. Maurice Swanson) were cultured in the same medium except FBS was increased to 20%. All cells were kept in a humidified incubator with 5% CO_2_ at 37°C.

For Nano-Trap experiments involving Neuro2a, cells were transfected with Transporter 5 (Kyfora Bio, 26008) according to manufacturer’s instructions, then collected after overnight expression for immunoprecipitation. All other transfections were performed using Lipofectamine 2000 (Fisher Scientific, 11-668-019) with 24- or 40-hour expression for splicing analysis.

### Immunoprecipitation

Neuro2a cells were collected after transfection by scraping from 10cm^2^ plates with ice cold phosphate-buffered saline (PBS) and centrifuging at 500g in a 4°C centrifuge for 3 minutes. Removing the supernatant, the cell pellet was resuspended in ice cold PBS and again spun at 500g for 3 minutes; this process was then repeated one more time. After the washes, the cell pellets were resuspended with lysis buffer (10mM Tris-HCl (pH 7.5), 150mM NaCl, 0.5mM EDTA, 0.5% NP-40) containing protease/phosphatase inhibitor (Halt Protease and Phosphatase Inhibitor, Thermo Scientific, 78441). The resuspended pellet was incubated on ice for 30 minutes with extensive pipetting every 10 min. The lysate was spun at 17,000g in the 4°C centrifuge for 10 minutes and the supernatant collected. Protein concentration was determined using Pierce BCA Assay (Thermo Scientific, 23227), and 10μg of protein was set aside to be run as input. From the remaining supernatants, equal amounts of protein were incubated with 25μL of GFP-Trap (Proteintech, gtma-10) or RFP-Trap magnetic agarose (Proteintech, rtma-10), rotating for 1 hour at 4°C. The beads were washed three times with wash buffer (10mM Tris-HCl (pH 7.5), 150mM NaCl, 0.5mM EDTA, 0.05% NP-40) and boiled in 2x Laemmli buffer (Bio-Rad, 1610737) at 95 °C for 10 minutes, unless noted otherwise. The Laemmli buffer was supplemented with reducing agent β-Mercaptoethanol (β-ME) or Dithiothreitol (DTT) as indicated. After boiling, the beads were magnetically separated and the sample left behind was used for western blot analysis.

For nuclear-cytoplasmic fractionation or IPs performed on the nuclear fraction of transfected cells, the cell pellet was first resuspended in lysis buffer (20mM Tris-HCl (pH 7.5), 10mM KCl, 2mM MgCl_2_, 1mM EGTA, 0.1% NP-40) and centrifuged at 1,000 g for 5 minutes at 4°C. The supernatant can be saved as the cytoplasmic fraction, while the remaining pellet is washed with wash buffer (20mM Tris-HCl (pH 7.5), 150mM KCl, 2mM MgCl_2_, 1mM EGTA, 0.2% NP-40) and centrifuged again at 1,000 g for 3 minutes in the refrigerated centrifuge. The resulting pellet, containing nuclei, was resuspended with DNase buffer (20mM Tris-HCl (pH 7.5), 100mM NaCl, 2mM MgCl_2_, 1mM CaCl_2_, 1% Triton X-100), supplemented with endonuclease (Benzonase, EMD Millipore, 70664-3, 1:200), and incubated on ice for 20-30 minutes. After centrifugation at 1,000 g for 3 minutes at 4°C, the nuclei were lysed with RIPA buffer (25mM Tris-HCl (pH 7.5), 150mM NaCl, 0.1% SDS, 0.5% sodium deoxycholate, 1% Triton X-100) for 20-30 minutes on ice, after which the lysate was spun at 2000 g for 3 minutes at 4°C and the supernatant was collected. All lysis, wash, and DNase buffers contained protease/phosphatase inhibitor. Protein concentration was again determined using Pierce BCA Assay and equal amounts of protein were incubated with 25 μL RFP-Trap or GFP-Trap magnetic agarose rotating for 1 hour at 4°C. The beads were washed three times with RIPA buffer and boiled in 2x Laemmli buffer at 95 °C for 10 minutes.

Immunoprecipitation of endogenous MBNL1 and MBNL2 was conducted using a modified CLIP protocol (37). Neuro2a cells, plated the day before, were scraped off with ice-cold PBS and centrifuged at 500 g for 4 min. With the supernatant removed, each pellet was resuspended in 500 uL lysis buffer (50mM Tris-HCl (pH 7.5), 100mM NaCl, 1mM MgCl_2_, 0.1mM CaCl_2_, 1% NP-40, 0.1% sodium deoxycholate, 0.05% SDS), containing protease/phosphatase inhibitor and endonuclease. For IPs involving human fibroblasts, Benzonase was replaced with Micrococcal Nuclease (Thermo Scientific, 88216), which required the addition of CaCl_2_ to a final concentration of 5mM. The resuspended pellet was incubated on ice for 30 min with extensive pipetting every 10 min. During this time, 100 uL of protein A Dynabeads (Invitrogen, 10001D) or Sheep anti-Mouse IgG Dynabeads (Invitrogen, 11031) were transferred to a 1.5 mL microcentrifuge tube and washed 3X with lysis buffer. After the third wash, the beads were resuspended in 200 uL of lysis buffer with 3 ug of rabbit α-MBNL1 antibody (Millipore Sigma, ABE241) or normal Rabbit IgG (Millipore Sigma, 12–370) added for the MBNL1 IP or 5μg of mouse α-MBNL2 3B4 antibody (Santa Cruz, sc-136167) or normal Mouse IgG (Millipore Sigma, 12–371) added for the MBNL2 IP. The antibody-bead mixture was then rotated at 4 °C for 30–60 min. Meanwhile, the cell lysates were centrifuged at maximum speed for 3 min, the supernatant was carefully collected, and the protein concentration was determined using Pierce BCA Assay. The antibody-conjugated beads were washed again 3X with lysis buffer and an equal amount of protein was added to each. Samples were rotated overnight at 4 °C. The next day, the beads were washed two times with high-salt wash buffer (50mM Tris-HCl (pH 7.5), 1M NaCl, 1mM EDTA, 1% NP-40, 0.1% sodium deoxycholate) and boiled in 2x Laemmli buffer, containing or lacking β-Mercaptoethanol, for 5 minutes at 95°C. After boiling, the beads were magnetically separated and the eluted sample used for western blot analysis.

For IPs involving mouse brain, the tissue was harvested from E17.5 and adult C57/BL6 mice, weighed, and suspended in 10 volumes of lysis buffer, including protease/phosphatase inhibitor and Benzonase, per gram of tissue. The tissue was disrupted with a Dounce homogenizers and incubated on ice for 10 minutes. The lysates were cleared by centrifugation at 4°C for 15 minutes at 7,500 g and the supernatant used for endogenous immunoprecipitation described above.

### Western Blotting

Immunoprecipitated proteins were separated on 4–20% Tris-Glycine polyacrylamide gradient gels (Bio-Rad, 4561096) and transferred onto 0.45 mm nitrocellulose membranes. Membranes were blocked in 5% BSA in PBS before being incubated overnight at 4 °C with primary antibody diluted in 5% BSA in PBS containing 0.1% Tween 20 (PBS-T). The following day, membranes were incubated with IRDye 680LT donkey anti-mouse (1:20000, LICOR, 926-68072) and either IRDye 680LT donkey anti-rabbit (1:20000, LICOR, 926-68073) or HRP mouse anti-rabbit IgG light chain (1:5000, Jackson ImmunoResearch, 211-032-171), diluted in 5% BSA in PBS-T, for 1 hour at RT. All blots were also incubated with Rhodamine-conjugated α-GAPDH human Fab fragment (1:1000, Bio-Rad, 12004168). HRP was detected using Tanon™ High-sig ECL Western Blotting Substrate (ABclonal, 180-5001) and the blots were imaged on the ChemiDoc MP system and quantified using Image Lab software (Bio-Rad). Any quantification of signal from western blot bands included normalization to GAPDH housekeeping protein.

### RNA sequencing and Differential Splicing Analysis

RNA was extracted transfected from WT and DKO MEFs using the Quick-RNA Miniprep Kit (Zymo Research, R1054) with on-column DNase I digestion. Total RNA was Poly-A enriched by Novogene and sequenced on a paired end 150bp cycle. Each sample had at least >22 million reads. Raw reads were trimmed with fastp (version 0.23.4) using default settings. Trimmed reads were then aligned with STAR (version 2.7.11a) using a Gencode M33 149 bp built index with default settings. BAM outputs from the STAR alignment were then used to perform differential splicing analysis using rMats-Turbo (version 4.2), with the following settings: --readlength 150, --variable-read-length, --allowclipping. To visualize splicing differences, inclusion levels were obtained from the rMats junction output files and plotted in R (version 4.3.3) ggplot2 (version 3.5). Quantification of EGFP was done using Salmon (version 1.10.2), by mapping trimmed reads with the default settings to a mm39 decoy-aware index containing GRCm39 cDNA and ncRNA sequences (v110) along with the EGFP sequence from the MBNL1 constructs. MEF MBNL1/2 HITS-CLIP wig files were obtained from GEO: GSE60487. Using BEDOPS (version 2.4.41) wig files were converted to bed format using the -wig2bed command. The final bed file containing HITS-CLIP sites were converted to mm39 coordinates using the UCSC genome browser liftover tool.

### Reverse Transcription PCR (RT-PCR)

RNA was extracted from transfected MEFs and human fibroblasts using either the Quick-RNA Miniprep Kit (Zymo Research, R1054) or the Direct-zol RNA Miniprep Kit (Zymo Research, R2050) with the on-column DNase I digestion. RNA was reverse transcribed using the High-Capacity cDNA Reverse Transcription kit (Applied Biosystems, 4368814) with random hexamers according to the manufacturer’s protocol. PCR was performed on the resulting cDNA using primers that spanned the segment of interest and complemented constitutive exons of the respective gene (*Tcea2* FWD: *tgccaagtccctcatcaagtc*, REV: *tcacggatggcctccttagt*; *Wnk1* FWD: *agtcagcctcaagtgtcagc*, REV: *tttgttctggaggagcagcc*). Primers for exon 4 of the *TNNT2* mini-gene (RTB300) and *MBNL1* exon 7 have been reported previously (28). All cDNA was PCR-amplified 30 cycles, except RTB300 which was amplified 28 cycles, using Hot Start *Taq* 2x Master Mix (New England Biolabs, M0496S). The *Wnk1* products were run on a 1% TBE agarose gel with SYBR Safe (Invitrogen, S33102), the *Tcea2* products on a precast 1% TBE agarose gel with ethidium bromide (Bio-Rad, 1613028), and the RTB300 and MBNL1 products on precast 3% TBE agarose gels with ethidium bromide (Bio-Rad, 1613030). All gels were run for 1 hour at 100V then imaged on the ChemiDoc MP system and quantified using Image Lab software.

### Generation of full-length and truncated GFP-TCEA2

The presence or absence of intron 6 in *Tcea2* was verified by Sanger sequencing (Genewiz, USA) of the gel purified RT-PCR products. We created primers that span the coding sequence of *Tcea2* which lacks intron 6, and results in a full-length version of the protein, and *Tcea2* that possesses intron 6, which contains a premature termination codon. The primers were also designed to generate Acc65I and BamHI restriction enzyme sites for insertion into pEGFP-C1 vector. The resulting GFP-TCEA2 plasmids were confirmed by Plasmidsaurus using Oxford Nanopore Technology with custom analysis and annotation.

### Immunofluorescence (IF)

Transfected Neuro2a were washed once with PBS, fixed with 4% paraformaldehyde (PFA) diluted in PBS for 10 min, then washed 3×5 min with PBS. Cells were permeabilized with 0.2% Triton X-100 in PBS before being washed again 3×5 min with PBS. Neuro2a transfected with GFP-MBNL1/2 isoforms and mutants or mCherry-3xFlag-MBNL1-41 and the Ex. 7,8,10 fragment were mounted with ProLong Glass containing NucBlue (Invitrogen, P36981). N2A transfected with GFP-TCEA2 plasmids were incubated with HCS CellMask Deep Red Stain (Fisher Scientific, H32721) diluted in PBS (1:10000) for 10 min before 3×5 min PBS washes and mounting. All steps were carried out at room temperature (RT).

### RNA Fluorescence *In Situ* Hybridization (FISH)

Before starting, all prepared solutions were made fresh using diethyl pyrocarbonate treated water and RNase-free reagents. First, cells were fixed with 4% PFA for 10 min at RT and washed 3× with 1X PBS. Cells were then washed once with 2X saline-sodium citrate (SSC) for 10 minutes at RT and once with 2X SSC containing 10% formamide, which was pre-warmed to 37°C, for 5 minutes at 37°C. Cells were pre-hybridized with hybridization buffer (10% formamide, 2X SCC, 0.05X PBS, 10% Dextran Sulfate, 10mM Ribonucleoside Vanadyl Complexes, 2μg/μL BSA, 0.66μg/μL Salmon Sperm DNA, 0.66μg/μl tRNA) for 1.5 hours in a 37°C humidified chamber. Afterwards, cells were transferred to hybridization buffer containing CAG_10_ Cy3 (IDT) at a concentration of 1.85μM and again incubated at 37°C in a humidified chamber for 16 hours. After hybridization, the coverslips were washed twice for 20 minutes each at 37°C in SSC/10% formamide that had been pre-warmed to 37°C. Next, the cells were washed with 2X SSC 3 times successively then 2 times for 5 minutes each at RT. For FISH experiments including IF, the cells were immediately incubated in IF blocking buffer for 1 hour at RT then with primary antibody in IF blocking buffer overnight at 4°C. After washing 3× with PBS, secondary antibody incubation in IF blocking buffer was performed for 1 hour at RT with the following: donkey α-mouse Alexa 488 (1:500, Jackson ImmunoResearch, 715-545-150). The coverslips were washed 3 more times with PBS and mounted with ProLong Glass containing NucBlue.

### Fixed Cell Imaging

Fixed cell imaging of transfected Neuro2A cells and DKO MEFs and human fibroblasts was performed using widefield illumination on a Nikon Eclipse Ti-E inverted fluorescent microscope with a Plan-Neofluar 1.4 NA x60 oil objective. 15-20 images were taken in a z-series at 2 μm steps and deconvolved using a 3-D blind constrained iterative algorithm (AutoQuant, CyberMetrics). All images are displayed as maximum intensity projections of the deconvolved slices.

### Image Analysis

Imaris imaging software (Bitplane) was used to count foci and measure their volume in DKO MEFs co-transfected with wiltype or C325A GFP-MBNL1-41 and DMPKS or 480CTG plasmids. Discreet spots that contained colocalization of CUG RNA signal and GFP-MBNL1 signal were deemed legitimate foci for the purpose of counting. For volume measurement, the ‘Coloc’ module of Imaris was used to create a new channel based on colocalized pixels. 3-D volumes were created from this colocalization channel, which were used to calculated total foci volume and average foci volume.

### Statistics and Reproducibility

All experiments were performed at least three times. Images shown in panels represent consistent results observed across all replicates. Statistical analyses performed are referenced in each figure legend. All analyses were performed using GraphPad Prism software (Version 10.0.2).

## RESULTS

### Endogenous MBNL1 forms reducing agent-sensitive high molecular weight species in cell lines and embryonic mouse brain tissues

Previous work has demonstrated that immunoprecipitation (IP) of GFP-MBNL1 and elution without boiling produces a novel western blot band located at twice the expected molecular weight of MBNL1, representing an MBNL1 dimer (27). To verify that endogenous MBNL1 forms a dimer, we immunoprecipitated (IPed) MBNL1 from mouse (Neuro2a) and human neuroblastoma cell lines (SH-SY5Y), eluting with sample buffer containing or lacking β-ME. In the sample eluted with buffer lacking β-ME, we observed a MBNL1 band twice its expected molecular weight, which was absent in the sample eluted with buffer containing β-ME (**Supplementary Figure 1A**). The dimer band was also absent from the input lanes, indicating that a limited proportion of MBNL1 exists as a dimer (**Supplementary Figure 1A**). We also performed an IP of endogenous MBNL1 from embryonic (E17.5) and adult mouse brain to demonstrate that MBNL1 dimerization also occurs in tissue. A MBNL1 dimer band, as well as a band located at a molecular weight three times that of MBNL1 monomer, was observed in the sample IPed from E17.5 mouse brain and eluted with sample buffer lacking β-ME (**Supplementary Figure 1B**). Notably, these high molecular weight (HMW) bands were greatly reduced in MBNL1 that was IPed from adult mouse brain and eluted under non-reducing conditions (**Supplementary Figure 1B**). Comparing the intensities of both HMW bands and the monomeric band, we found that the MBNL1 HMW-to-monomer ratio was significantly higher in embryonic brain compared to adult brain (**Supplementary Figure 1C**). Both HMW bands were also absent from all input lanes and MBNL1 IPed and eluted under reducing condition (**Supplementary Figure 1B**). Sensitivity to β-ME suggested that the MBNL1 dimer and HMW species are held together by one or more disulfide bond formed between cysteine residues located in MBNL1.

### A cysteine residue in exon 7 of MBNL1 is necessary for its homodimerization via disulfide bond formation

Given the sensitivity of the MBNL1 dimer to reducing conditions and the established connection between MBNL1 exon 7 and dimerization (26,27), we hypothesized that one or more cysteine residues in MBNL1 exon 7 form intermolecular disulfide bonds. Exon 7 is located in the unstructured C-terminal domain of MBNL1, which also contains a bipartite NLS that spans exons 5 and 6 (**Figure 1A**). Upon inspection, we found a cysteine in exon 7 that is conserved among MBNL orthologues (**Figure 1A**). To test its involvement in MBNL1 homodimerization, we mutated the exon 7 cysteine (Cys325) to alanine in GFP tagged MBNL1-41, a 41kDa MBNL1 isoform that lacks exon 5 and is present in both the nucleus and cytoplasm (**Figure 1B**). We used GFP tagged MBNL-40, which lacks both exon 5 and exon 7, as a negative control (**Figure 1B**). Mutation of Cys325 to alanine (C325A) did not appear to alter the distribution or expression level of GFP-MBNL1-41 in Neuro2a cells (**Supplementary Figure 2A**). In parallel to imaging, we performed GFP IP on transfected Neuro2a cells, again eluting under either reducing or non-reducing conditions. A band corresponding to a GFP-MBNL1 dimer was only present in the lane containing GFP-MBNL1-41 that was IPed and eluted with sample buffer lacking β-ME (**Figure 1C**). The dimer band was absent in samples of IPed GFP-MBNL1-41 eluted with β-ME and samples of IPed GFP-MBNL1-40 and GFP-MBNL1-41 C325A eluted without β-ME (**Figure 1C**). Like the endogenous MBNL1 IPs, the dimer band was absent from all input lanes, suggesting that, even when overexpressed, very little MBNL1 exists as a dimer. To verify that β-ME is not the only reducing agent capable of breaking the disulfide bond holding the MBNL1 dimer together, we tested another commonly used reducing agent, DTT. When added to elution buffer, DTT eliminated the dimer band in IPed GFP-MBNL1-41 in a concentration dependent manner (**Supplementary Figure 2D**).

**Figure 1.**
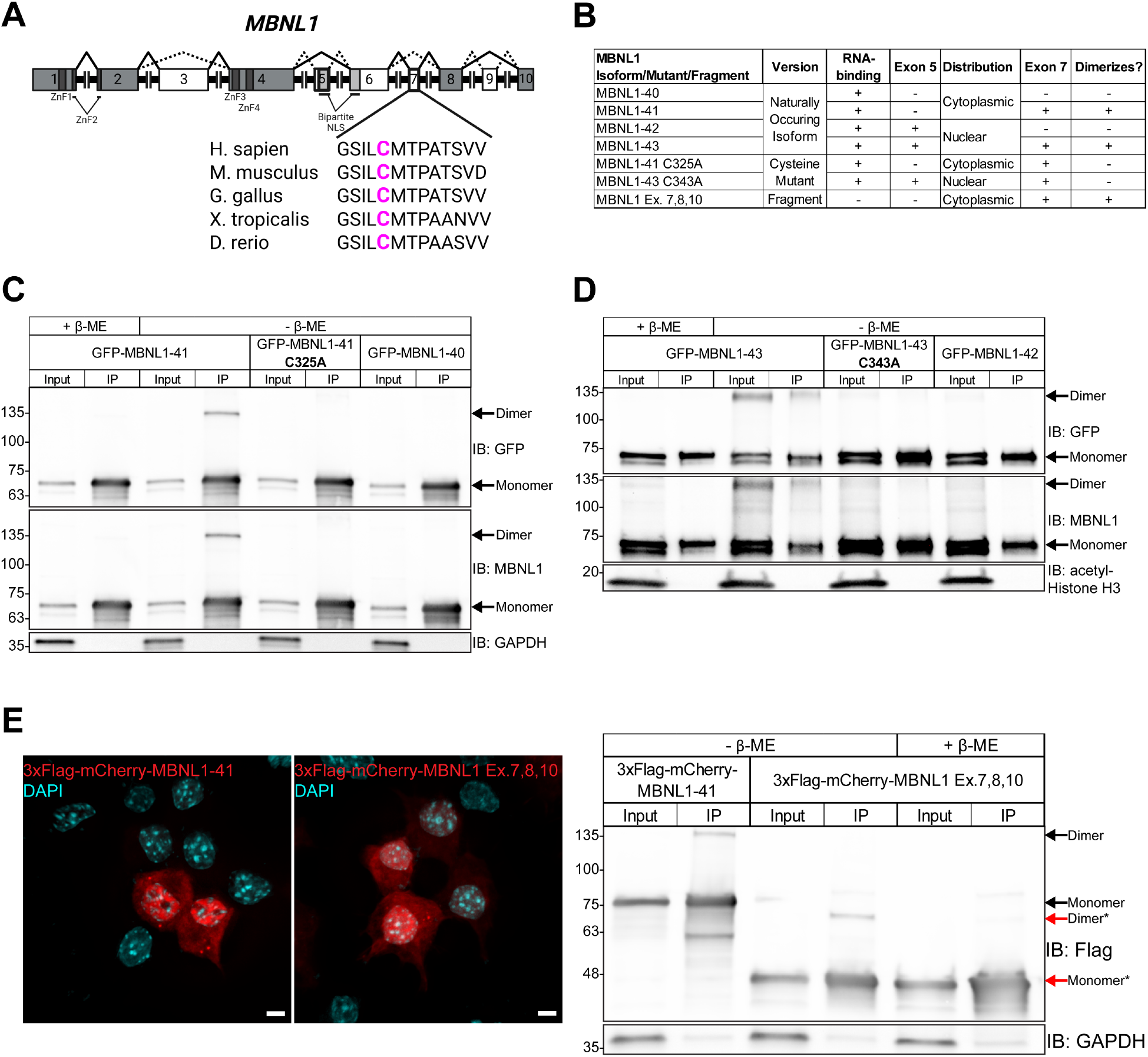
A cysteine residue in exon 7 of MBNL1 is responsible for its homodimerization. (**A**) Exon and protein domain structure of human MBNL1. Constitutive exons are colored gray and alternative exons white. The canonical splicing pattern is represented by solid lines, while alternative patterns are represented by dashed lines. A callout of exon 7 shows the location of a cysteine residue conserved among multiple species. Created in BioRender. Bassell Lab, G. (2025) https://BioRender.com/h81i289. (**B**) Table listing MBNL1 isoforms, mutants, and fragments used throughout Figure 1. The presence of exon 5, which determines nucleocytoplasmic distribution, and exon 7, which determines capacity for dimerization, are indicated for each protein. (**C**) Representative western blot against GFP, MBNL1, and GAPDH following IP of GFP-MBNL1-40, -41, and -41 C325A from transfected Neuro2a cells. Samples were prepared or eluted with Laemmli buffer containing or lacking β-ME. (**D**) Representative western blot against GFP, MBNL1, and acetylated Histone H3 following IP of GFP-MBNL1-42, -43, and -43 C343A from nuclear lysate of transfected Neuro2a cells. Samples were prepared or eluted with or without β-ME. (**E**) (Left) Representative images showing the distribution of full length 3xFlag-mCherry-MBNL1-41 and a 3xFlag-mCherry-tagged fragment of MBNL1 Exons 7, 8, 10 in transfected Neuro2a cells. Scale bars = 5µm. (Right) Representative western blot against Flag and GAPDH following IP of full-length and fragment versions of 3xFlag-mCherry-MBNL1. Samples were prepared or eluted with or without β-ME. *indicates species composed of the 3xFlag-mCherry-MBNL1 Exon 7,8,10 fragment

Next, we tested whether nuclear MBNL1 isoforms, MBNL1-42 and MBNL1-43, which contain the full bipartite NLS, are also able to form dimers (**Figure 1B,D**). The exon 7 cysteine in GFP-MBNL1-43, Cys343, was again mutated to alanine (C343A) (**Figure 1B**), and GFP-MBNL1-42, which lacks exon 7, was used as a negative control. Imaging revealed no major differences in expression or distribution between wildtype and mutant GFP-MBNL1-43 (**Supplementary Figure 2B**). We performed another GFP IP on the nuclear fraction of transfected Neuro2a cells and found that the dimer band was now found not only in the lane containing GFP-MBNL1-43 that was IPed and eluted without β-ME, but also in the corresponding input lane (**Figure 1D**). These results suggest that the amount of MBNL1 present as a dimer is higher in the nucleus than in the cytosol, and that this high molecular weight band is not an artifact of IP. We ruled out the influence an N-terminal GFP tag might have on MBNL dimerization by swapping the GFP for a HA tag and moving it to the C-terminus of MBNL1-43 and MBNL1-43 C343A (**Supplementary Figure 2C**). The fact that IP was unnecessary for visualization of the more abundant nuclear dimer allowed us to probe other properties of the MBNL1 dimer. Following fractionation of Neuro2a cells transfected with MBNL1-43-HA or MBNL1-43-HA C343A, we prepared the nuclear lysate with sample buffer containing β-ME, but varied the amount of time the lysate was boiled at 95°C. Western blot of these samples revealed that the MBNL1-43-HA dimer band was present when the samples were not boiled, but it gradually disappeared with increasing boiling time (**Supplementary Figure 2C**). However, in line with our findings, the dimer band was absent in the MBNL1-43-HA C343A sample even without boiling (**Supplementary Figure 2C**). The greater abundance of MBNL1 dimerization in the nucleus intimated a possible nuclear function for the MBNL1 dimer.

We included RNase in all IPs conducted for MBNL dimerization to rule out high molecular weight species that might form due to multiple proteins binding the same RNA transcript. However, to test whether RNA-binding is a prerequisite for MBNL1 dimerization, we employed a fragment of MBNL1 that lacks exons 1-6, rendering it unable to bind RNA (**Figure 1E**). Both full length MBNL1-41 and this C-terminal fragment containing exon 7 were fused to 3xFlag-mCherry, allowing us to perform an RFP IP, elute with sample buffer lacking or containing β-ME, and compare dimerization. Bands twice the respective expected molecular weights of full length and fragmented 3xFlag-mCherry-MBNL1 were present in the samples IPed and eluted with sample buffer lacking β-ME (**Figure 1E**). The 3xFlag-mCherry-MBNL1 fragment dimer band disappeared when β-ME was included in the IP elution buffer (**Figure 1E**). The results of this IP demonstrate that RNA-binding is not a prerequisite for dimerization, and that exons 7, 8, and 10, contain the minimum residues sufficient for MBNL1 dimerization.

### Endogenous MBNL2 also forms reducing agent-sensitive dimers in a process that is developmentally regulated

Exon 7 is also conserved in MBNL2, a paralog of MBNL1, but no research has been conducted into its dimerization. We also detected a high molecular weight band in samples of endogenous MBNL2, that were IPed from Neuro2a or SH-SY5Y cells and eluted in sample buffer lacking β-ME (**Supplementary Figure 3A**). This high molecular weight band, which was twice the expected molecular of MBNL2, is the first evidence supporting the existence of an MBNL2 dimer. Like MBNL1, the MBNL2 dimer band was absent in samples eluted in reducing conditions (**Supplementary Figure 3A**). Sensitivity to reducing conditions suggesting that the MBNL2 dimer is also held together by at least one disulfide bond. Additionally, the MBNL2 dimer band was missing from the input lanes, indicating a scarcity similar to that of the MBNL1 dimer (**Supplementary Figure 3A**). We also IPed MBNL2 from embryonic (E17.5) and adult mouse brain. A dimer band was observed for MBNL2 that was IPed from E17.5 mouse brain and eluted with sample buffer lacking β-ME (**Supplementary Figure 3B**). Like the IPs conducted on MBNL1, the dimer band was absent from input lanes and MBNL2 IPed and eluted under reducing condition (**Supplementary Figure 3B**). However, unlike MBNL1, a dimer band was also detected in the lane with MBNL2 that was IPed from adult mouse brain (**Supplementary Figure 3B**). This might be due to MBNL2 expression being higher than MBNL1 in adult brain tissue (38,39). Nevertheless, the ratio of MBNL2 dimer to monomer differed greatly between E17.5 and adult mouse brain. A strong MBNL2 dimer band in embryonic brain decreased slightly in adult brain, whereas monomeric MBNL2, which was quite weak in embryonic brain, intensified significantly in adult brain (**Supplementary Figure 3B**). This resulted in a significantly greater MBNL2 dimer-to-monomer ratio in embryonic brain compared to adult brain (**Supplementary Figure 3C**). This suggests that, while MBNL2 levels increase during developmental of the brain (**Supplementary Figure 3C**), the predominant form of MBNL2 shifts from dimeric to monomeric.

### Multiple cysteine residues in the C-terminus contribute to MBNL2 dimerization

Like MBNL1, the endogenous MBNL2 dimer was also sensitive to reducing conditions, leading us to hypothesize that it also contains at least one cysteine residue capable of forming an intermolecular disulfide bond. Three major isoforms of MBNL2 exist in humans: Isoform 1 (MBNL2-I1), Isoform 3 (MBNL2-I3), and Isoform 4 (MBNL2-I4). Apart from exons 5, 9, and 10, the other seven exons of MBNL1 are all conserved in MBNL2. Unique to MBNL2, however, is its final C-terminal exon, which is alternatively spliced to yield either a short, 3 amino acid-long tail, or a long, 41 amino acid-long tail, the latter of which includes 4 cysteine residues exclusive to MBNL2 (**Figure 2A,B**). MBNL2-I1 possesses the conserved exon 7 from MBNL1 and the short tail, making it functionally equivalent to MBNL1 from the standpoint of dimerization (**Figure 2A,B,C**). MBNL2-I3 and MBNL2-I4, on the other hand, both possess the long tail unique to MBNL2 (**Figure 2A,B**). We began by confirming that the properties of cysteine residues do not change in the context of the MBNL2 C-terminal domain. To do this, we mutated the single, conserved cysteine (Cys324) in a GFP-tagged version of MBNL2-I1 to alanine (C324A) (**Figure 2C**). When we performed a GFP IP, a band corresponding to a GFP-MBNL2 dimer was only present in the lane containing wildtype GFP-MBNL2-I1 that was IPed and eluted with sample buffer lacking β-ME (**Figure 2C**). This GFP-MBNL2 dimer band disappeared when β-ME was included in the elution buffer (**Figure 2C**). The GFP-MBNL2 dimer was absent from IPed GFP-MBNL2-I1 C324A, even when β-ME was excluded from the elution buffer (**Figure 2C**). The C324A mutation did not appear to affect the distribution or expression of GFP-MBNL2-I1 when transfected and imaged in Neuro2a cells (**Supplementary Figure 4A**). We proceeded to perform GFP IPs on GFP tagged versions of the remaining two MBNL2 isoforms to assess whether the cysteines unique to MBNL2 support dimerization and whether varying number of cysteines influence the amount of MBNL2 present as a dimer (**Figure 2D**). However, we first had to consider a recent study, which revealed a PEST degradation sequence located in the long version of the MBNL2 tail, resulting in variable protein levels among MBNL2 isoforms (40). Quantifying MBNL2 signal from the input lanes of our GFP IPs, we were able to corroborate findings from the study: normalized GFP-MBNL2-I1 signal was significantly greater than GFP-MBNL2-I3 and GFP-MBNL2-I4 (**Figure 2E**). However, despite differences in protein levels, all three GFP-MBNL2 isoforms formed dimer bands when IPed and eluted with sample buffer lacking β-ME, resulting in similar dimer-to-monomer ratios (**Figure 2E**). When β-ME was included in the elution buffer for GFP-MBNL2-I4, which contains the largest number of cysteines, the dimer band disappeared (**Figure 2D**). The existence of multiple cysteine residues unique to MBNL2 led us to wonder whether all are required for MBNL2 dimerization.

**Figure 2.**
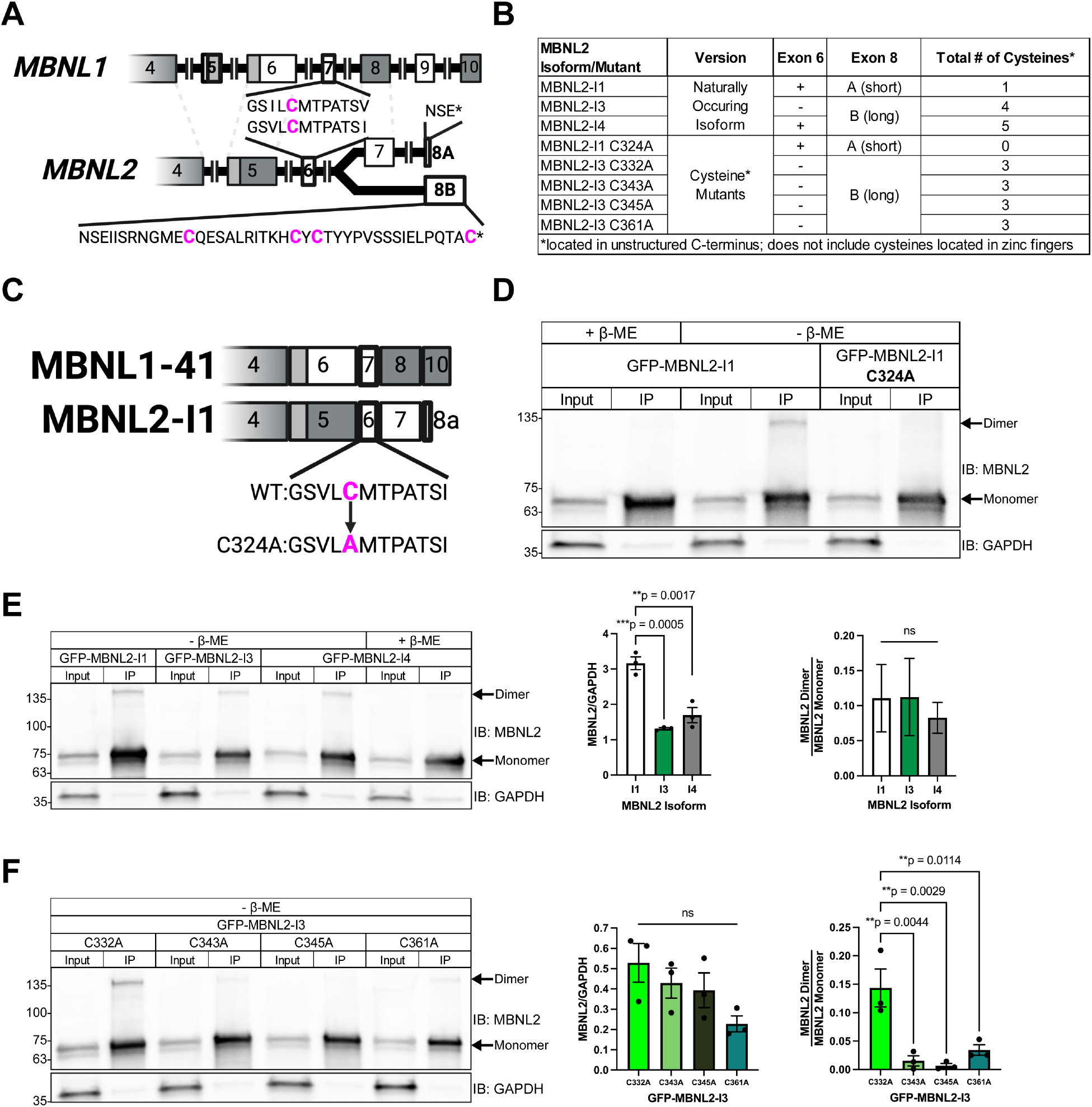
The carboxy-terminal domain of MBNL2 contains multiple cysteines that allow it to homodimerize. Comparison of exonic structure between the C-terminal domains of *MBNL1* and *MBNL2*. Constitutive exons are colored gray and alternative exons white. Dotted lines indicate conservation of exons between *MBNL1* and *MBNL2*. Callouts show the location of the cysteine residues within conserved exon 7/6 and the unique, long form of exon 8 (B) in MBNL2. Created in BioRender. Bassell Lab, G. (2025) https://BioRender.com/y89o916. (**B**) Table listing MBNL2 isoforms and mutants used in Figure 2. The presence of exon 6 and either the short (A) or long form of exon 8 are indicated for each protein, along with the total number of C-terminal domain cysteines. (**C**) Illustration of the C-terminal domains of MBNL1-41 and MBNL2-I1 along with the C324A mutation in the latter. Created in BioRender. Bassell Lab, G. (2025) https://BioRender.com/y89o916. (**D**) Representative western blot against MBNL2 and GAPDH following IP of GFP-MBNL2-I1 and GFP-MBNL2-I1 C324A from transfected Neuro2a cells. Samples were prepared or eluted with Laemmli buffer containing or lacking β-ME. (**E**) (Left) Representative western blot against MBNL2 and GAPDH following IP of GFP-MBNL2-I1, -I3, and -I4 from transfected Neuro2a cells. Samples were prepared or eluted with or without β-ME. (Center) Quantification of GFP-MBNL2 isoform levels normalized to GAPDH and (Right) dimer-to-monomer ratio of IPed GFP-MBNL2 isoforms. Mean and standard error are reported across 3 biological replicates. *\*\*p*<0.01, *\*\*\*p*<0.001 by one-way ANOVA with Tukey’s post-hoc test. (**E**) (Left) Representative western blot against MBNL2 and GAPDH following IP of GFP-MBNL2-I3 C332A, C343A, C345A, and C361A from transfected Neuro2a cells. All samples were prepared or eluted in Laemmli buffer lacking β-ME. (Center) Quantification of each GFP-MBNL2-I3 mutant normalized to GAPDH and (Right) dimer-to-monomer ratio of IPed GFP-MBNL2-I3 mutants. Mean and standard error are reported across 3 biological replicates. *\*\*p*<0.01 by one-way ANOVA with Tukey’s post-hoc test.

As we had already confirmed that changing the single conserved cysteine in MBNL2-I1 abolishes its dimerization, we decided to focus solely on the cysteine residues unique to MBNL2. We serially mutated each of the four cysteine residues in the C-terminus of GFP-MBNL2-I3, which lacks the conserved exon 7 from MBNL1 and thus contains only cysteines exclusive to MBNL2 (**Figure 2B**). This resulted in four GFP-MBNL2-I3 mutants: GFP-MBNL2-I3 C332A, GFP-MBNL2-I3 C343A, GFP-MBNL2-I3 C345A, and GFP-MBNL2-I3 C361A (**Figure 2B**). When we performed GFP IPs on these mutants and eluted without β-ME, only MBNL2-I3 C332A formed a dimer comparable to unmutated GFP-MBNL2-I3 (**Figure 2E,F**). Mutation of any of the remaining single cysteine residues abolished or greatly reduced the amount of GFP-MBNL2-I3 present as a dimer (**Figure 2F**). None of the mutations affected GFP-MBNL2-I3 distribution or expression based on imaging performed on transfected Neuro2a cells (**Supplementary Figure 4B**). These results indicate Cys332 is dispensable for MBNL2-I3 dimerization, yet each of the other cysteines is necessary for dimerization.

### RNAseq conducted on MBNL1/2 DKO MEFs expressing WT or C325A GFP-MBNL1-41 reveals novel splicing events regulated by MBNL1 dimerization

The higher rate of MBNL1 dimerization in the nucleus pointed to a possible nuclear function for the MBNL1 dimer. Among the nuclear functions of MBNL1, the most highly studied is its regulation of alternative splicing (15-17). To get a clearer picture of how dimerization might affect MBNL1-dependent alternative splicing regulation, we moved to a system free of endogenous MBNL, namely fibroblasts derived from *Mbnl1*^*-/-*^; *Mbnl2*^*-/-*^ mouse embryos (DKO MEFs) (18,31). We confirmed, by western blot, that WT and C325A GFP-MBNL1 could be re-introduced in DKO MEFs by transient transfection (**Figure 3A,B**). Just as in Neuro2a cells, a dimer band was present when WT GFP-MBNL1-41 was IPed from transfected DKO MEFs and eluted with sample buffer lacking β-ME (**Figure 3A**). Additionally, the dimer band was only detected in the IP lane of DKO MEFs expressing GFP-MBNL1-41; it was absent from all input lanes and the IP lane of DKO MEFs expressing GFP-MBNL1-41 C325A (**Figure 3A**). Levels of both WT and C325A GFP-MBNL1-41 were also similar across biological replicates probed by western blot (**Figure 3B**), ensuring that any changes in splicing are due to dimerization and not differences in protein expression or stability. With this system in place, we moved forward with exploring a possible role for dimerization in MBNL1-mediated alternative splicing regulation.

**Figure 3.**
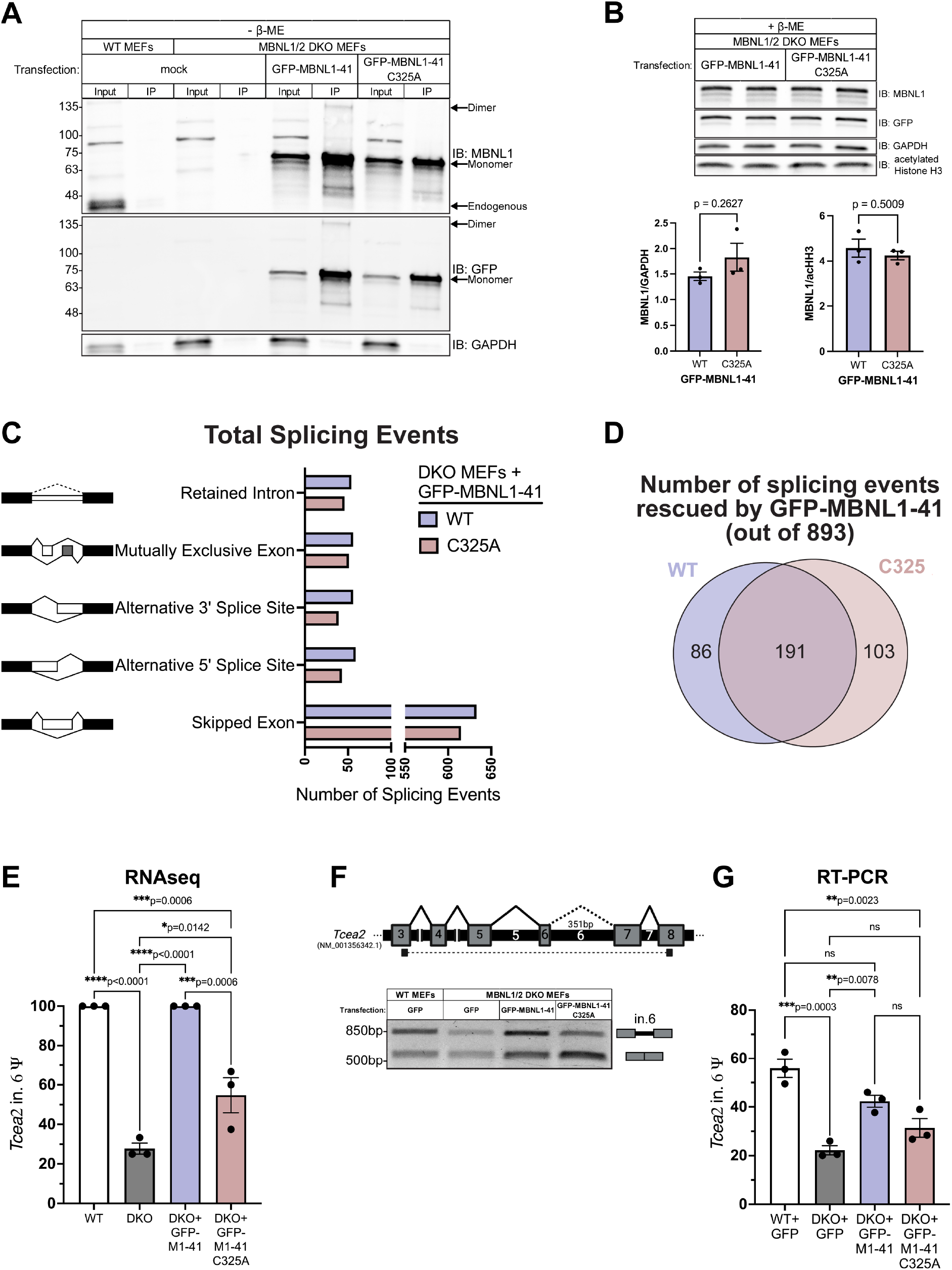
RNAseq conducted on MBNL1/2 DKO MEFs expressing WT or C325A GFP-MBNL1-41 reveals novel splicing events affected by MBNL1 dimerization. (**A**) Representative western blot against MBNL1, GFP, and GAPDH following GFP IP from wildtype MEFs or MBNL1/2 DKO MEFs transfected with WT or C325A GFP-MBNL1-41 or mock transfected. All samples were prepared or eluted in Laemmli buffer lacking β-ME. (**B**) (Above) Representative western blot against MBNL1, GFP, GAPDH, and acetylated Histone H3 (acHH3) for whole cell lysate from DKO MEFs expressing either WT and C325A GFP-MBNL1-41. Samples were prepared with Laemmli buffer containing β-ME. (Below) Quantitation of MBNL1 signal normalized to GAPDH (left) or acHH3 (right). Mean and standard error are reported from 3 biological replicates. *p*>0.05 by two-tailed unpaired t test. (**C**) All types of alternative splicing are regulated by both WT and C325A MBNL1-41. (**D**) Venn diagram of aberrant splicing events in DKO MEFs that are rescued to WT MEF levels (|ΔΨ|≤0.1) by expression of WT, C324A, or both GFP-MBNL1-41 species. (**E**) Psi values for intron 6 of *Tcea2* from RNAseq conducted on WT MEFs, DKO MEFs, and DKO MEFs transfected with either WT or C325A GFP-MBNL1-41. Data is reported as mean and standard error of 3 biological replicates. *\*p*<0.05, *\*\*\*p*<0.001, *p\*\*\*\**<0.0001 by one-way ANOVA with Tukey’s post-hoc test. (**F**) (Above) Schematic depicting the location of primers used to measure *Tcea2* intron 6 Ψ by RT-PCR. Created in BioRender. Bassell Lab, G. (2025) https://BioRender.com/i05v148. (Below) Representative agarose gel showing *Tcea2* splicing products generated by RT-PCR on RNA from WT MEFs, DKO MEFs, and DKO MEFs expressing either WT or C325A GFP-MBNL1-41. (**G**) Quantitation of *Tcea2* intron 6 Ψ values from RT-PCR splicing gels in **F**. Data is reported as mean and standard error of 3 biological replicates. *\*\*p*<0.01, *\*\*\*p*<0.001 by one-way ANOVA with Tukey’s post-hoc test.

To accomplish this, we began by performing RNAseq on three independent wild type MEF clones and three independent DKO MEF clones to establish endogenous targets of MBNL. We also included a separate set of three DKO MEF clones transfected with either GFP-MBNL1-41 or GFP-MBNL1-41 C325A. The RNAseq reads were mapped by STAR (41) to genome and splice junctions, yielding approximately 176.6M, 147.3M, 154M, and 158.15M uniquely mapping reads from the three sets of WT MEFs, DKO MEFs, and DKO MEFs reconstituted with WT or C325A GFP-MBNL1-41, respectively. Ψ values were quantitated across all alternative splicing events (ASEs) using rMATS-turbo (42). A similar number of all types of ASEs, skipped cassette exons, retained introns, mutually exclusive exons and alternative 5′ or 3′ splice sites, were regulated by both WT and C325A GFP-MBNL1-41 when overexpressed in DKO MEFs (**Figure 3C**). Of the 893 ASEs significantly altered between WT and DKO MEFs (FDR<0.05 and |ΔΨ|≥0.1), 191 were rescued (|ΔΨ|≤0.1) by both WT and C325A GFP-MBNL1-41, 86 were rescued exclusively by WT GFP-MBNL1-41, and 103 were unexpectedly rescued only by C325A GFP-MBNL1-41 (**Figure 3D**).

Among the 86 ASEs rescued by reconstitution with WT GFP-MBNL1-41 alone, retention of intron 6 in Transcription Elongation Factor A2 (*Tcea2*) showed the greatest rescue and least variability (**Figure 3E**). MBNL-dependent intron retention has not been studied extensively, so, to ensure the veracity of this result, we designed primers spanning *Tcea2* intron 6 and performed RT-PCR (**Figure 3F**). After resolving the RT-PCR products and measuring the isoform ratios, we were able to validate the results of our RNAseq: intron 6 was significantly excluded from *Tcea2* in DKO MEFs, reconstitution with the WT GFP-MBNL1-41 resulted in a partial, but significant, rescue, and reconstitution with C325A GFP-MBNL1-41 failed to rescue *Tcea2* intron 6 retention (**Figure 3G**). We also submitted the RT-PCR products for Sanger sequencing, which confirmed the top band did indeed contain intron 6 (**Supplementary Figure 5A**). While more subtle in appearance, a similar pattern was observed when DKO MEFs were reconstituted with WT and C343A GFP-MBNL1-43 (**Supplementary Figure 5B**). Upon closer inspection of the Sanger sequencing chromatogram, we noticed an in-frame, premature termination codon (PTC) located within intron 6 of *Tcea2* (**Supplementary Figure 5C**). While this would normally result in non-sense mediated decay (43), the presence of a discernible band on agarose gels suggested otherwise. If translated, retention of intron 6 would result in a distinct 32 amino acid-long tail and premature truncation of the TCEA2 protein (**Supplementary Figure 5C**). We decided to clone the full-length, canonical *Tcea2* coding sequence and the *Tcea2* coding sequence containing this intronic PTC into a GFP vector (**Supplementary Figure 5D**). Both full-length and truncated GFP-TCEA2 were able to be overexpressed in Neuro2a cells and resolved by SDS-PAGE. Western blot bands for both proteins showed similar intensity and were located at their expected molecular weights (**Supplementary Figure 5D**). Given the role of TCEA2 in transcription initiation and preventing R-loop formation (44-46), the data above suggest dimerization does play a role in MBNL1-dependent regulation of an ASE with potentially important downstream consequences.

### MBNL1 dosage-dependent splicing events can also be affected by the ability of MBNL1 to dimerize

Given only 380 of 893 MBNL-dependent ASEs were fully rescued to WT MEF levels (|ΔΨ|≤0.1) by reconstitution of either WT or C325A GFP-MBNL1-41 in DKO MEFs, we wanted to examine the effect on the remaining 513 ASEs that did not reach the threshold of full rescue. For both fully rescued and subthreshold events, we plotted their average ΔΨ from WT MEFs when DKO MEFs were expressing either WT GFP-MBNL1-41, measured on the x-axis, or C325A GFP-MBNL1-41, measured on the y-axis (**Figure 4A**). The large number of MBNL-dependent ASEs that were not fully rescued, as evidenced by their absence from colored zones representing |ΔΨ|≤0.1 (**Figure 4A**), led us to wonder whether this was due to variability in expression of GFP-MBNL1-41 between or even within replicates. Using EGFP Transcripts Per Million (TPM) as a readout of GFP-MBNL1-41 expression, we verified that expression levels of WT and C325A GFP-MBNL1-41 were similar within each replicate (**Figure 4B**). However, the expression level between replicates was quite variable (**Figure 4B**). This was unintended but could be due to several factors including the number of times the DKO MEFs had been passaged, the density of the DKO MEFs, and the number of freeze-thaw cycles the plasmids had gone through. Nevertheless, we exploited this variability in expression which enabled us to analyze the RNAseq results on a more granular level and identify ASEs that are not only sensitive to MBNL1 dimerization, but also sensitive to MBNL1 concentration.

**Figure 4.**
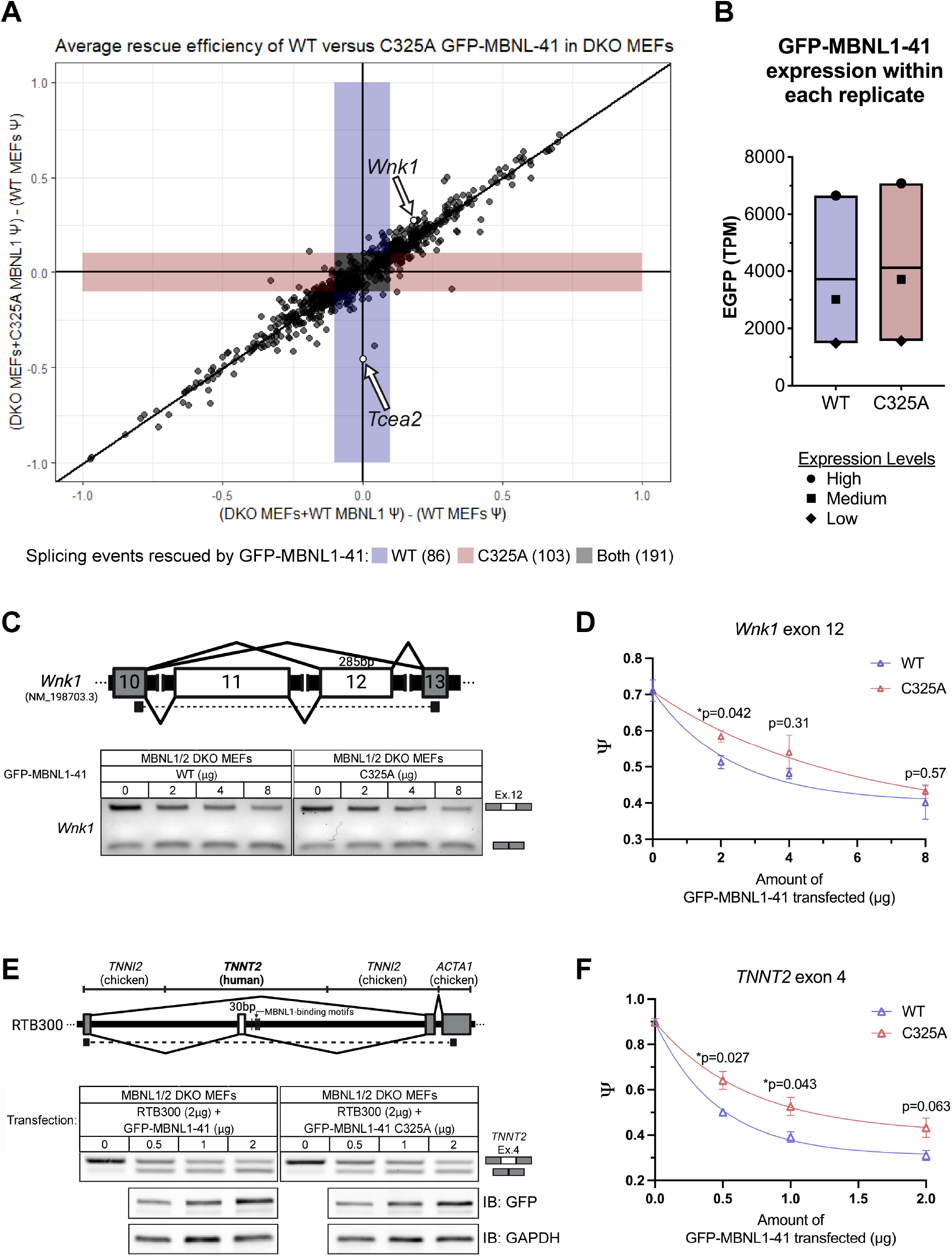
Splicing events affected by MBNL1 dimerization can also be sensitive to MBNL1 concentration. A dot plot comparing the average ΔΨ from WT MEF levels for 893 aberrant splicing events in MBNL1/2 DKO MEFs reconstituted with either WT or C325A GFP-MBNL1-41. This includes events that are fully rescued (|ΔΨ|≤0.1) by WT (blue), C325A (red), or both (grey) GFP-MBNL1-41 species. All other events (white) fall below the threshold of full rescue. Average ΔΨ values were calculated from the 3 biological replicates submitted for RNAseq. Retention of *Tcea2* intron 6, which is rescued only by WT GFP-MBNL1-41 (see **Figure 3**), is labeled. Expression levels of WT and C325A GFP-MBNL1-41 for each replicate submitted for RNAseq, measured as TPM of EGFP. Each replicate is represented by different symbol based on its GFP-MBNL1-41 expression level, labeled “High”, “Medium”, or “Low”. (**C**) (Above) Schematic depicting the location of *Wnk1* exon 12 RT-PCR primers; constitutive exons are gray and alternative exons white. Created in BioRender. Bassell Lab, G. (2025) https://BioRender.com/i05v148. (Below) Representative splicing gel of DKO MEFs transfected with varying amounts of WT or C325A GFP-MBNL1-41. (**D**) MBNL1 WT/C325A dose-response curves for *Wnk1* exon 12 splicing. Psi values are plotted against the amount of WT (blue) or C325A (red) GFP-MBNL1-41 transfected, and a one-phase decay curve fitted to the resulting data points. Means and standard errors were calculated from 3 biological replicates. Statistical significance between WT and C325A was calculated for each amount of GFP-MBNL1-41 using two-tailed unpaired t tests, *\*p*<0.05. (**E**) (Above) Schematic depicting the location of RT-PCR primers for *TNNT2* minigene (RTB300); the gene and species of each segment of the mini-gene is listed above. Created in BioRender. Bassell Lab, G. (2025) https://BioRender.com/i05v148. (Below) Representative RT-PCR splicing gels of DKO MEFs co-transfected with RTB300 and varying amounts of WT or C325A GFP-MBNL1-41. Representative western blots of the corresponding protein samples are shown underneath. (**F**) MBNL1 WT/C325A dose-response curves for *TNNT2* exon 4 splicing. Psi values are plotted against the amount of WT (blue) or C325A (red) GFP-MBNL1-41 transfected, and a one-phase decay curve fitted to the resulting data points. Means and standard errors were calculated from 3 biological replicates. Statistical significance between WT and C325A was calculated for each amount of GFP-MBNL1-41 using two-tailed unpaired t tests, *\*p*<0.05.

Concentration dependent behavior has been attributed to multiple, known MBNL1-dependent ASEs (47). A closer review of the 513 sub-threshold ASEs revealed numerous splicing events which were rescued by WT but not C325A GFP-MBNL1-41 only in the “low” expression replicate. These same ASEs showed no difference between WT and C325A GFP-MBNL1-41 in the “medium” and “high” expression replicates. For example, skipping of exon 12 in Lysine Deficient Protein Kinase 1 (*Wnk1*) is a previously established MBNL-dependent ASE (17), which is among the sub-threshold ASEs that skew towards partial rescue by WT GFP-MBNL1-41 in the dot plot of the average of all three RNAseq replicates (**Figure 4A**). After closer examination of *Wnk1* exon 12 Ψ values in the DKO MEF replicates expressing “high”, “medium”, and “low” levels of WT and C325A GFP-MBNL1-41, it became apparent that *Wnk1* exon 12 skipping is both dosage-and dimerization-dependent (**Supplementary Figure 6**). For the “low” level replicates, *Wnk1* exon 12 ΔΨ from WT MEF levels was 0.074 when WT GFP-MBNL1-41 was expressed in DKO MEFs but 0.553 when C325A GFP-MBNL1-41 was expressed (**Supplementary Figure 6A**,**E**). However, in the “medium” replicates, *Wnk1* exon 12 |ΔΨ| was 0.325 and 0.211 when WT and C325A GFP-MBNL1-41 were expressed, respectively (**Supplementary Figure 6B**,**E**), and in the “high” replicates, *Wnk1* exon 12 |ΔΨ| was 0.153 and 0.102 for WT and C325A GFP-MBNL1-41, respectively (**Supplementary Figure 6C**,**E**). With primers designed to span *Wnk1* exon 12, we performed RT-PCR on RNA isolated from DKO MEFs transfected with varying amounts of WT or C325A GFP-MBNL1-41 (**Figure 4C**). We were able to produce a similar MBNL1 dosage- and dimerization-dependent pattern for *Wnk1* exon 12 Ψ (**Figure 4D**). Analysing Ψ for *Wnk1* exon 12 at different GFP-MBNL1-41 expression levels allowed us to determine that *Wnk1* exon 12 skipping is a MBNL1 dimerization-dependent ASE, but only when exposed to specific concentrations of MBNL1. Other sub-threshold ASEs are likely sensitive to MBNL1 dimerization, but this result is confounded at higher expression levels, where MBNL that is unable to dimerize is functionally interchangeable with MBNL that is able to dimerize.

Another well-established MBNL1-dependent splicing event, skipping of exon 4 in the cardiac muscle Troponin T (*TNNT2*) gene (15), exhibits different dose-dependent responses when MBNL1 contains or lacks exon 7 (28). The mouse ortholog of this human gene was not detected in WT or DKO MEFs so, to test if this difference is due to MBNL1 dimerization, we co-transfected DKO MEFs with the established *TNNT2* mini-gene RTB300 (36) and varying amounts of WT or C325A GFP-MBNL1-41 (**Figure 4E**). We were able to verify that GFP-MBNL1-41 C325A elicits a dose-dependent response similar to that elicited by GFP-MBNL1-40, namely it promotes exclusion of *TNNT2* exon 4 less efficiently than GFP-MBNL1-41, especially at lower concentrations (**Figure 4F**) (28). Successful replication of a MBNL1 dosage and exon 7-dependent ASE, not found in our RNAseq dataset, using GFP-MBNL1-41 C325A, further bolstered our conclusion that MBNL1 dimerization via disulfide bond plays an important role in its regulation of alternative splicing.

### MBNL1 forms higher order species in DM1 patient fibroblasts

Despite the prominent role MBNL plays in DM1, no consideration has been given to the potential impact of MBNL dimerization on DM1 pathology and vice versa. Aberrant inclusion of *Mbnl1* exon 7 *has* been observed in muscle from a mouse model of DM1 (35) and human DM1 patients (28), but the outcome of this aberrant inclusion has not been pursued further at the level of protein. To study the impact DM1 has on MBNL1 dimerization, we turned to fibroblasts from individuals with congenital DM1, which possess over 1700 CTG repeats compared to 5 CTG repeats in control fibroblasts. The DM1 fibroblasts, which possess both higher levels of nuclear MBNL1 when fractionated and MBNL1/2 positive CUG foci when imaged (**Supplementary Figure 7A**,**B**), also had higher levels of *MBNL1* exon 7 inclusion (**Supplementary Figure 7D**). CUG foci and associated nuclear sequestration of MBNL1 or MBNL2 were absent in control fibroblasts (**Supplementary Figure 7A**,**C**), which also lacked exaggerated *MBNL1* exon 7 inclusion (**Supplementary Figure 7D**). IP of MBNL1 from WT and DM1 fibroblast samples revealed a dimer band in both when eluted with Laemmli buffer lacking β-ME (**Supplementary Figure 7E**). However, an additional band at a molecular weight greater than that of the dimer was present only in DM1 samples eluted without β-ME (**Supplementary Figure 7E**). While human MBNL1 possesses only one cysteine residue capable of forming a disulfide bond, this finding implies MBNL1 might form a heterodimer with MBNL2 or another cysteine-containing protein in DM1. MBNL1 and MBNL2 are both sequestered to CUG foci, and MBNL2 isoforms contain multiple cysteines that could form disulfide bonds with more than one other MBNL molecule. While this result reveals a potential impact of DM1 on MBNL1/2 multimerization, we also sought to investigate the inverse: the impact of MBNL1 dimerization on DM1.

### MBNL1 dimerization affects the number and size of CUG repeat foci

We continued using DKO MEFs to decipher potential contributions of MBNL1 dimerization to DM1 pathogenesis; its lack of endogenous MBNL and our ability to overexpress specific MBNL1 species make it an ideal system. However, first we wanted to confirm that the C325A mutation does not affect nuclear sequestration of MBNL1 to expanded CUG foci. Neuro2a cells were co-transfected with WT or C325A GFP-MBNL1-41 and a plasmid containing a fragment of human *DMPK* with 0 (DMPKS) or 480CTG repeats, the latter of which is meant to recapitulate DM1 pathogenesis. When co-transfected with 480CTG repeats, GFP-MBNL1-41 C325A signal was significantly increased in the nuclear fraction, similar to when WT GFP-MBNL1-41 was expressed (**Figure 5A,B**). Despite its increase in the nucleus due to sequestration, GFP-MBNL1-41 C325A still did not produce a dimer band in the absence of β-ME (data not shown), prompting us to investigate the appearance of the RNA foci in the presence of GFP-MBNL1-41 C325A.

**Figure 5.**
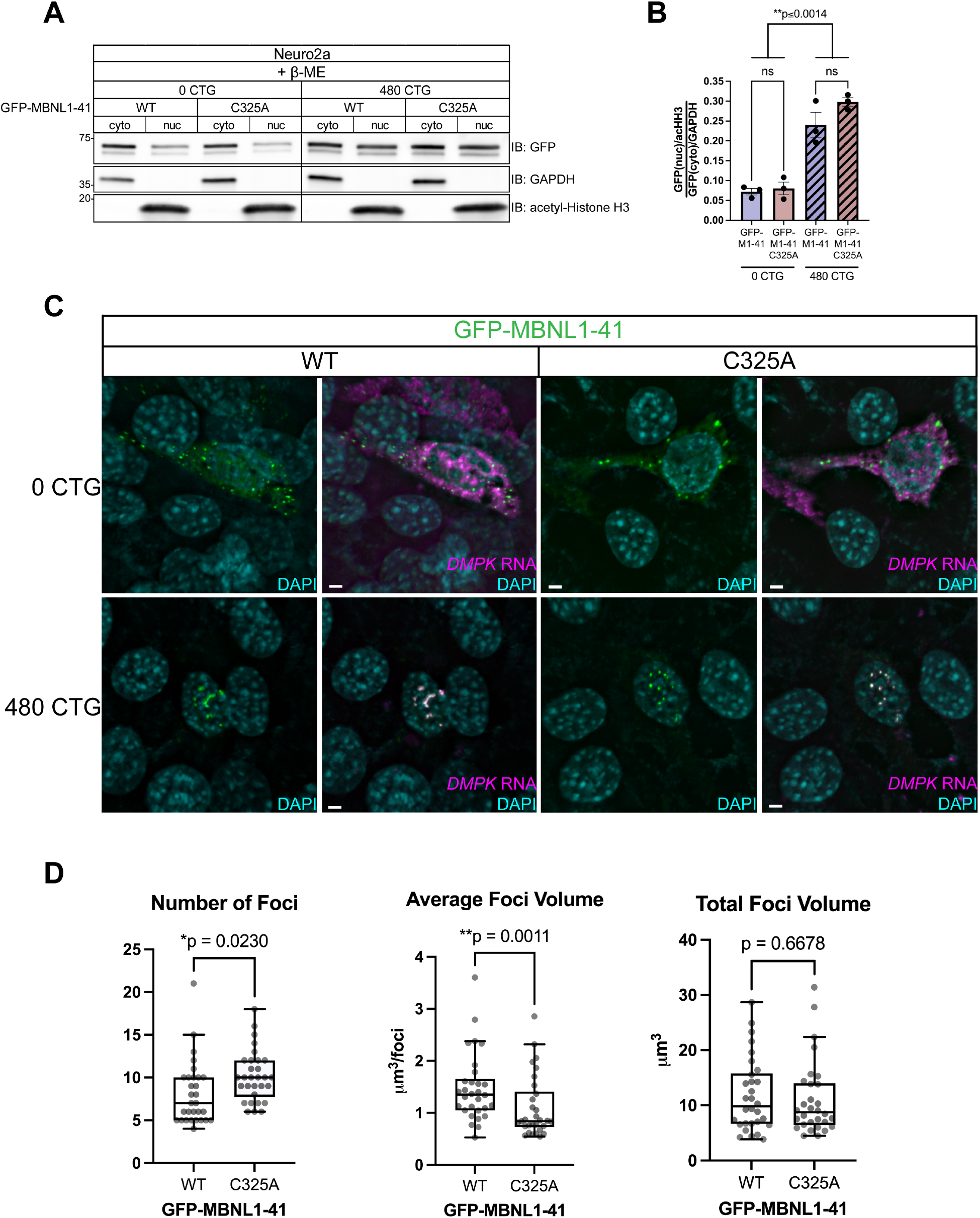
MBNL1 dimerization is necessary for maintenance of DM1 foci integrity. (**A**) Representative western blot following nucleocytoplasmic fractionation of Neuro2a cells co-transfected with WT or C325A GFP-MBNL1-41 and DMPKS or 480CTG repeats. GAPDH shows enrichment in the cytoplasmic fraction and acetylated Histone H3 in the nuclear fraction. All samples were prepared with Laemmli buffer containing β-ME. (**B**) Quantitation of nucleocytoplasmic ratio of GFP signal normalized to GAPDH (cytoplasmic) and acHH3 (nuclear) for Neuro2a co-transfected with WT or C325A GFP-MBNL1-41 and DMPKS or 480CTG repeats. Mean and standard error are reported from 3 biological replicates. *\*\*p*<0.01 by one-way ANOVA with Tukey’s post-hoc test. (**C**) Representative FISH images for *DMPK* RNA (magenta) in DKO MEFs co-transfected with WT or C325A GFP-MBNL1-41 and DMPKS or 480CTG plasmids. Scale bars = 3μm. (**D**) Quantitation of number of foci, average foci volume, and total foci volume in DKO MEFs co-transfected with 480CTG repeats and either WT or C325A GFP-MBNL1-41; n=30 cells across 3 biological replicates. Bars represent median, box outlines represent upper and lower quartiles, whiskers represent 10th and 90th percentiles. Two-tailed unpaired t test was conducted on number of foci and total foci volume. Non-parametric Kolmogorov-Smirnov test was conducted on average foci volume due to non-normal distribution of GFP-MBNL1-41 C325A samples. *\*p*<0.05, *\*\*p*<0.01

We ensured that the DKO MEFs could be co-transfected with GFP-MBNL1-41 and either the DMPKS or 480CTG repeat plasmids. The co-transfection was successful as evidenced by distinct nuclear foci positive for human *DMPK* RNA when FISH was conducted on DKO MEFs co-transfected with 480CTG repeats and either WT or C325A GFP-MBNL1-41 (**Figure 5C**). In contrast, when the DMPKS plasmid was co-transfected with either WT or C325A GFP-MBNL1-41, *DMPK* RNA FISH signal was dispersed in both the nuclear and cytoplasmic compartments (**Figure 5C**). Both WT and C325A GFP-MBNL1-41 colocalized entirely with the nuclear *DMPK* FISH signal in the DKO MEFs expressing 480CTG repeats but were mostly present in the cytoplasm as small granules in DKO MEFs expressing DMPKS (**Figure 5C**). While both WT and C325A GFP-MBNL1-41 were able to form foci when co-transfected with 480CTG repeats, the foci appeared smaller and more numerous when C325A GFP-MBNL1-41 was expressed versus when WT GFP-MBNL1-41 was expressed (**Figure 5C**). After quantifying the number and 3-dimensional volume of the foci, we found a significant difference in both number of foci (median foci number for WT vs. C325A: 7 vs. 10) and average foci volume (median average foci volume for WT vs. C325A: 1.353µm^3^/foci vs. 0.8428µm^3^/foci) per cell depending on the version of GFP-MBNL1-41 co-transfected with 480CTG repeats (**Figure 5D**). The total volume of foci per cell, on the other hand, was not significantly different between C325A and WT GFP-MBNL1-41 (**Figure 5D**). This result suggests that MBNL1 dimerization impacts the integrity of the toxic foci present in DM1.

## DISCUSSION

Previous evidence suggests that MBNL1 can self-associate through its exon 7 (26,27), but the nature of this interaction and its implications for RNA biology have remained unexplored. In this study, we identified a cysteine residue in exon 7 of MBNL1 (**Figure 1A**), which, when mutated to alanine (**Figure 1B**), eliminates a dimer band that normally forms when GFP-MBNL1-41 is IPed and eluted with sample buffer lacking β-ME (**Figure 1C**). Interestingly, immunoprecipitation was unnecessary to observe a dimer band in the nuclear fraction of Neuro2a expressing GFP-MBNL1-43 (**Figure 1D**), suggesting a potential nuclear function for the MBNL1 dimer. We also observed a high molecular weight band when endogenous MBNL2 was IPed and eluted without β-ME (**Supplementary Figure 3**); this is the first evidence supporting the existence of MBNL2 dimers. The C-termini of all three isoforms of human MBNL2 contain varying numbers of cysteine residues (**Figure 2A,B**), allowing each to dimerize when IPed as GFP-MBNL2 and eluted under non-reducing conditions (**Figure 2D**). Mutational analysis of cysteines unique to MBNL2 revealed that not all are necessary for its dimerization (**Figure 2E**). Given the greater proportion of MBNL1 present as a dimer in the nucleus, we pursued a potential function for MBNL1 dimerization in nucleus, namely in modulating MBNL1-dependent regulation of alternative splicing. By transfecting *Mbnl1*^*-/-*^;*Mbnl2*^*-/-*^ (DKO) MEFs with WT or C325A versions GFP-MBNL1-41 and performing RNAseq, we were able to identify novel ASEs affected by MBNL1 dimerization (**Figure 3**). Following careful analysis, we identified two classes of MBNL1-dimerization dependent ASEs: those that were sensitive to MBNL1 dimerization at all concentrations of MBNL1 and those that were sensitive to MBNL1 dimerization only at low concentrations of MBNL1 (**Figure 4**). After establishing a physiological role for MBNL1 dimerization, we explored a potential pathological role of MBNL1 dimerization. Despite the prominent role MBNL1 plays in DM1, MBNL1 dimerization has never been studied in the context of DM1. We imaged DKO MEFs co-transfected with WT or C325A GFP-MBNL1-41 and a fragment of *DMPK* containing 480 CTG repeats. We found that the CUG foci were smaller and more numerous when C325A GFP-MBNL1-41 was co-expressed versus when WT GFP-MBNL1-41 was co-expressed (**Figure 5C,D**). This result suggests that MBNL1 dimerization plays a role in maintaining the integrity of CUG foci in DM1. Overall, the findings in this study have revealed both a physiological and pathological role for MBNL1 dimerization dependent on disulfide bonds.

MBNL is by no means the only RBP that interacts with itself. In fact, during splicing regulation, most RBPs interact with each other, though not through disulfide bonds, but through non-covalent forces involving their IDRs (5). MBNL itself possesses an IDR in its C-terminal domain, which could theoretically allow it to dimerize through non-covalent interactions. However, to be clear, in this study we have uncovered functions unique to MBNL dimers that arise specifically through formation of disulfide bonds. If and how MBNL might dimerize through non-covalent forces between its unstructured C-terminal domain, and whether this is mutually exclusive from dimerization via disulfide bond formation fall outside the scope of the current study. MBNL is also not the only RBP that contains cysteine residues. An analysis of the amino acid composition of 1,555 RBPs showed that about 94% contain at least one cysteine residue (48, **Supplementary Figure 8**). Furthermore, among these 1,467 cysteine-containing RBPs, protein structure predictions revealed approximately 42% possess at least one cysteine located in a disordered region (**Supplementary Figure 8**). This is noteworthy because cysteine residues located in disordered regions are more likely to form disulfide bonds than cysteines located in highly ordered regions. One classic example of a RBP that dimerizes by forming a disulfide bond is Polypyrimidine Tract-Binding (PTB) protein (14). However, rather than splicing regulation, PTB de-dimerization was shown to enhance translation, specifically of *TP53* (14), a known RNA target of PTB (49). Based on evidence from this study, MBNL would represent one of the first RBPs whose self-interaction via a covalent, disulfide bond impacts its regulation of alternative splicing. However, the sheer number of RBPs that contain cysteine residues capable of forming disulfide bonds suggests there are likely other RBPs that form disulfide bonds as part of larger complexes that also regulate alternative splicing.

Not only do disulfide bonds serve as a means of connecting two cysteine-containing proteins, they also play an important role in the folding and stability of proteins. *Intra*molecular disulfide bonds can hold distinct portions of a protein together, maintaining the protein in a folded topology. This is especially relevant for unstructured regions of proteins, such as IDRs, where disulfide bonds can increase the orderedness of the region. However, disulfide bonds are also reversible, forming under oxidizing conditions and dissociating under reducing conditions (50). Because of this, disulfide bond-forming cysteines can promote both disorder-to-order and order-to-disorder transitions in IDRs depending on the redox conditions of the local environment (51). Changes in redox conditions can occur naturally or be caused by environmental stressors, such as reactive oxygen species. Redox-based conditional disorderedness of cysteine-containing IDRs can have important implications for the function of the larger protein (52,53). MBNL2 contains multiple cysteine residues in its unstructured C-terminal domain, which can form intermolecular disulfide bonds with other MBNL2 proteins, as demonstrated by this study. However, this does not rule out the possibility that intramolecular disulfide bonds can also form between these cysteine residues. When predictions of structural disorder are performed on MBNL2-I4 using the IUPred2A algorithm with Redox State parameters (54), its unstructured C-terminal domain does show significantly decreased disorderedness in oxidizing conditions (data not shown). This redox sensitivity is absent in MBNL1 and the other MBNL2 isoforms (data not shown), likely due to the smaller number of cysteine residues in their C-terminal domains. Further investigation is needed to determine the effect redox-dependent changes in disorderedness of MBNL2’s C-terminal domain might have on its function.

In this study we identified a novel MBNL1-dependent splicing events, retention of *Tcea2* intron 6, which is also sensitive to MBNL1 dimerization (**Figure 3D-G**). The TCEA2 protein plays two distinct, but equally important roles in the process of transcription, making any changes in its splicing pattern very consequential. During the early events of transcription, TCEA2 recruits the Mediator complex and RNA Polymerase II (RNA Pol II) to form the Preinitiation Complex at promoter sequences of protein-coding genes (44,45,55). Separately, TCEA2 also recognizes backtracked RNA Pol II during the elongation stage of transcription (56), stimulating transcript cleavage by RNA Pol II and resumption of transcription (57-59). When transcript cleavage is absent, the ability of backtracked RNA Pol II to resume transcription is greatly disturbed. This poses a problem for maintenance of genome stability because DNA-RNA hybrids called R-loops can form (46,56,60). There is a premature termination codon (PTC) present within *Tcea2* intron 6 which would normally lead to nonsense-mediated decay (NMD) when the intron is retained (43). However, a RT-PCR band corresponding to intron 6 inclusion and Sanger sequencing both confirmed that *Tcea2* transcripts containing intron 6 do exist in a nonnegligible amount (**Figure 3F & Supplementary Figure 5A**). Exceptions to NMD do exist, which are dictated by the location of the PTC and the length of the exon or intron in which the PTC is located (61). However, the PTC in *Tcea2* intron 6 does not fall under either of these exceptions (**Supplementary Figure 5C**). RBP binding motifs can also influence NMD efficiency (61); there are a couple MBNL1 motifs located in exon 7 downstream of intron 6, but it is unlikely that this is sufficient to affect NMD (**Supplementary Figure 5C**). It remains unclear how *Tcea2* transcripts containing intron 6 evade NMD and what effect this has on transcription and genome stability. In addition to *Tcea2* intron 6 retention, we found that skipping of exon 12 in *Wnk1*, a known MBNL1-dependent ASE (17), was also sensitive to MBNL1 dimerization but only at lower levels of MBNL1 expression (**Figure 4C-E**). This dovetails nicely with our finding that both MBNL1 and MBNL2 are expressed at low levels, but dimerize to a greater degree, in embryonic tissue (**Supplementary Figures 1B**,**C & 3B**,**C**). MBNL dimerization might play a greater role in regulating splicing transitions early in development, when MBNL expression is low, rather than later, when MBNL expression is higher. Dimerization could improve MBNL splicing efficiency to better utilize MBNL when it is in low amounts, but more extensive investigation would be needed to confirm this. Furthermore, exactly how dimerization improves MBNL splicing efficiency is unclear: it might change the affinity of MBNL for its target transcripts or even change the binding motif all together. Nevertheless, the precise mechanism explaining how MBNL1 dimerization regulates ASEs remains unknown.

There are several unanswered questions that arose from the above research, which fell outside the scope of this paper, but warrant continued investigation. One such question relates to the role that the multiple cysteine residues unique to MBNL2 play in its dimerization. Our data indicates that loss of any of the three C-terminal cysteines unique to MBNL2 greatly diminishes its ability to dimerize (**Figure 2F**). It is not clear why loss of a single cysteine residue nullifies the disulfide bond forming ability of the other two. Another interesting, albeit perplexing, finding from our RNAseq data was the existence of a sizable number of ASEs rescued exclusively by re-expression of GFP-MBNL1-41 C325A in DKO MEFs (**Figure 3D**). Our original hypothesis was that dimerization would facilitate most splicing events. It is rather likely that dimerization facilitates splicing in a dosage and context dependent manner. The monomeric form seems to be preferred at higher doses. It will also be important to know if disulfide bonds with other RBPs exist. Given the number of potential RBPs that could form disulfide bonds with MBNL1 (**Supplementary Figure 8**), future studies probing differences in the interactome between WT and C325A MBNL1 could reveal important insights into RBP and RNA biology. These are just some of the most outstanding questions that resulted from this research. However, despite the questions raised, overall, this research uncovered a novel instance of dimerization via disulfide bond formation involving an important RBP which has potentially major implications for RNA biology and disease pathogenesis.

## DATA AVAILABILITY

The RNAseq data is available at the Gene Expression Omnibus (GEO) under the accession number GSE291521.

## SUPPLEMENTARY DATA

Supplementary Data are available at NAR online.

## AUTHOR CONTRIBUTIONS

L.A.K., E.T.W., and G.J.B. conceptualization; A.K. data curation; L.A.K. and A.K. formal analysis; E.T.W. and G.J.B. funding acquisition; L.A.K. and E.X.Z. investigation; G.N.N. and G.J.B. project administration; K.R.M., L.S., R.P.H., E.T.W., and G.J.B. resources; A.K. software; E.T.W. and G.J.B. supervision; L.A.K., A.K., L.S., G.N.N., A.J.K., and G.J.B. visualization; L.A.K writing-original draft; L.A.K., E.T.W., and G.J.B. writing-review and editing

## ACKNOWLEDGEMENTS

We thank Dr. Maurice Swanson (University of Florida) for donating MBNL1/2 double knockout fibroblasts and congenital DM1 patient fibroblasts. We also thank Dr. Thomas Cooper (Baylor College of Medicine) for the plasmids carrying exons 11–15 of the *DMPK* gene expressing 0 (DMPKS) or 480 CTG repeats. Finally, we thank our colleagues Drs. Jie Jiang, Sulagna Das, Nisha Raj, and Zachary McEachin, as well as Dr. Karmella Haynes (Georgia Tech/Emory) and all members of the Bassell lab, for helpful discussion and insightful suggestions.

