## Supplemental_Figures for "Muscleblind-like proteins dimerize by forming disulfide bonds to regulate alternative splicing and pathogenic RNA foci formation"

#### Supplementary Figure 1

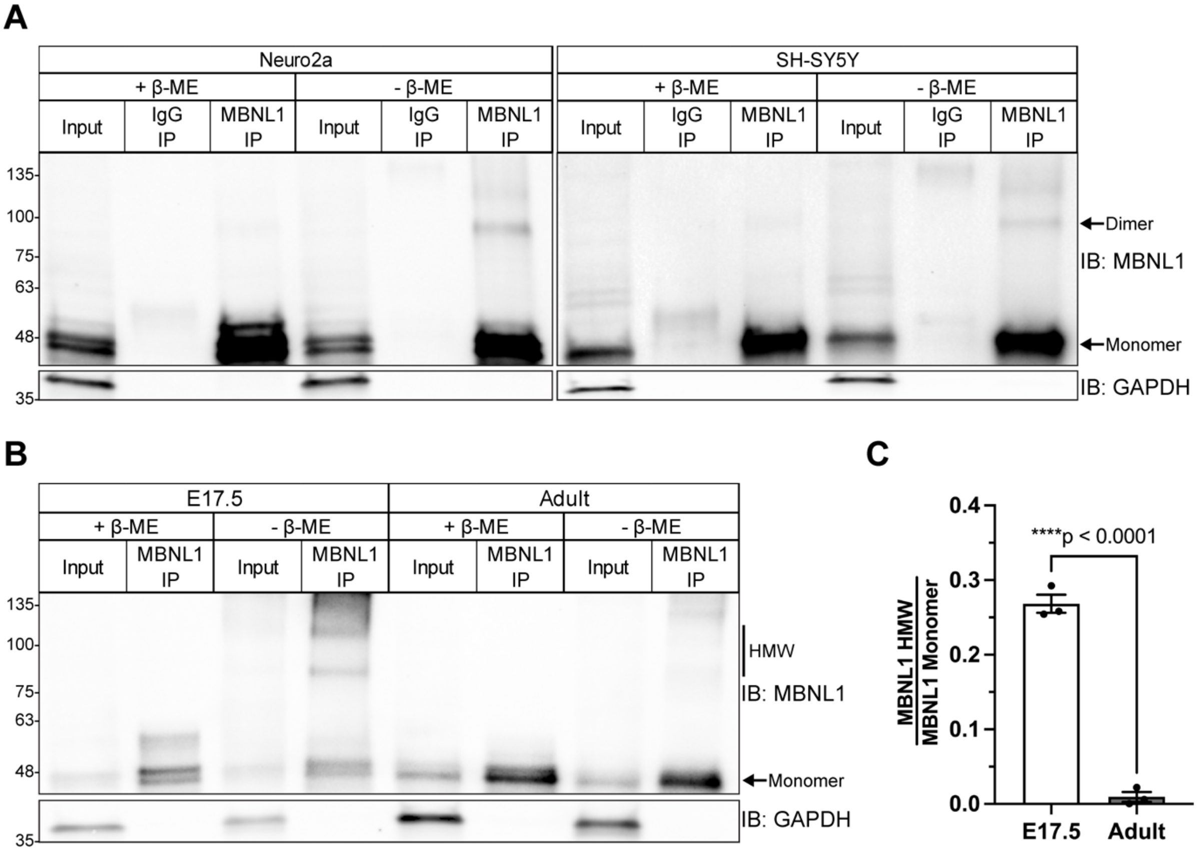


**Supplementary Figure 1.** MBNL1 forms dimers in mouse and human cell lines and high molecular weight species in embryonic mouse brain tissue. (**A**) Representative western blot of endogenous MBNL1 immunoprecipitated from Neuro2a and SH-SY5Y cells. Samples were prepared or eluted using Laemmli buffer containing or lacking β-mercaptoethanol (β-ME). (**B**) Representative western blot of endogenous MBNL1 immunoprecipitated from E17.5 and adult mouse whole brain lysate. Samples were prepared or eluted with Laemmli buffer lacking or containing β-ME. (**C**) Quantification of the dimer-to-monomer ratio of immunoprecipitated MBNL1 eluted without β-ME. Mean and standard error are reported from 3 biological replicates. ****$p$<0.0001 by two-tailed unpaired t test.

### Supplementary Figure 2

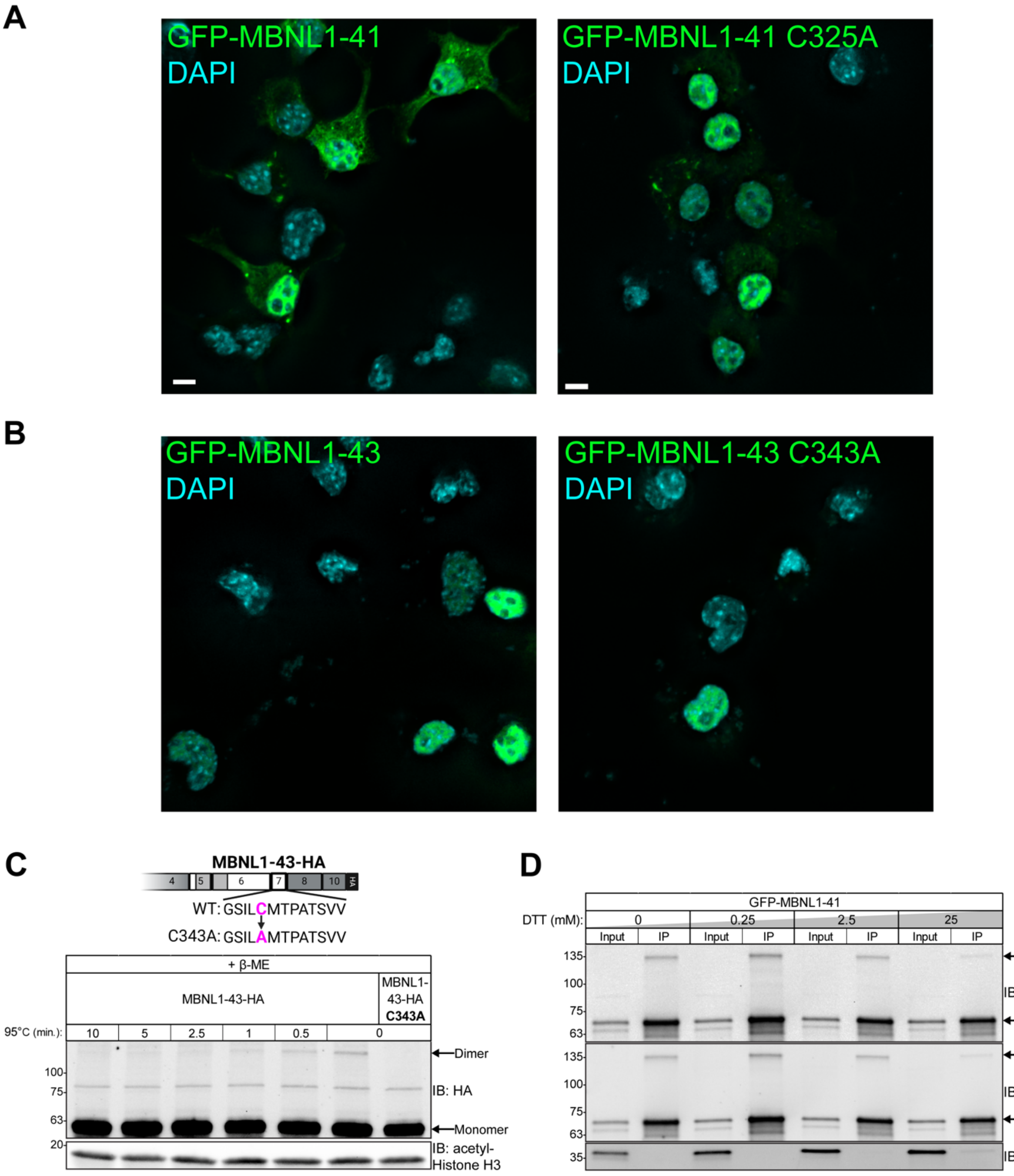


**Supplementary Figure 2.** Mutation of the cysteine in exon 7 of MBNL1 does not appear to alter its distribution, but both reducing agent and heat are required to break MBNL1 dimer.
(**A**) Representative images showing the distribution of GFP-MBNL1-41 and GFP-MBNL1-41 C325A in transfected Neuro2a cells. Scale bars = 5μm (**B**) Representative images showing the distribution of GFP-MBNL1-43 and GFP-MBNL1-43 C343A in transfected Neuro2a cells. (**C**) (Above) Diagram of the C-terminus of MBNL1-43-HA along with the C343A mutation. Created in BioRender. Bassell Lab, G. (2025) https://BioRender.com/h81i289. (Below) Representative western blot of the nuclear fraction of Neuro2a cells transfected with MBNL1-43-HA or MBNL1-

43-HA C343A. Samples were prepared with β-ME and boiled at 95°C for varying amounts of time. (**D**) Representative western blot against MBNL1, GFP, and GAPDH following immunoprecipitation of GFP-MBNL1-41, prepared or eluted with varying concentrations of Dithiothreitol (DTT).

#### Supplementary Figure 3

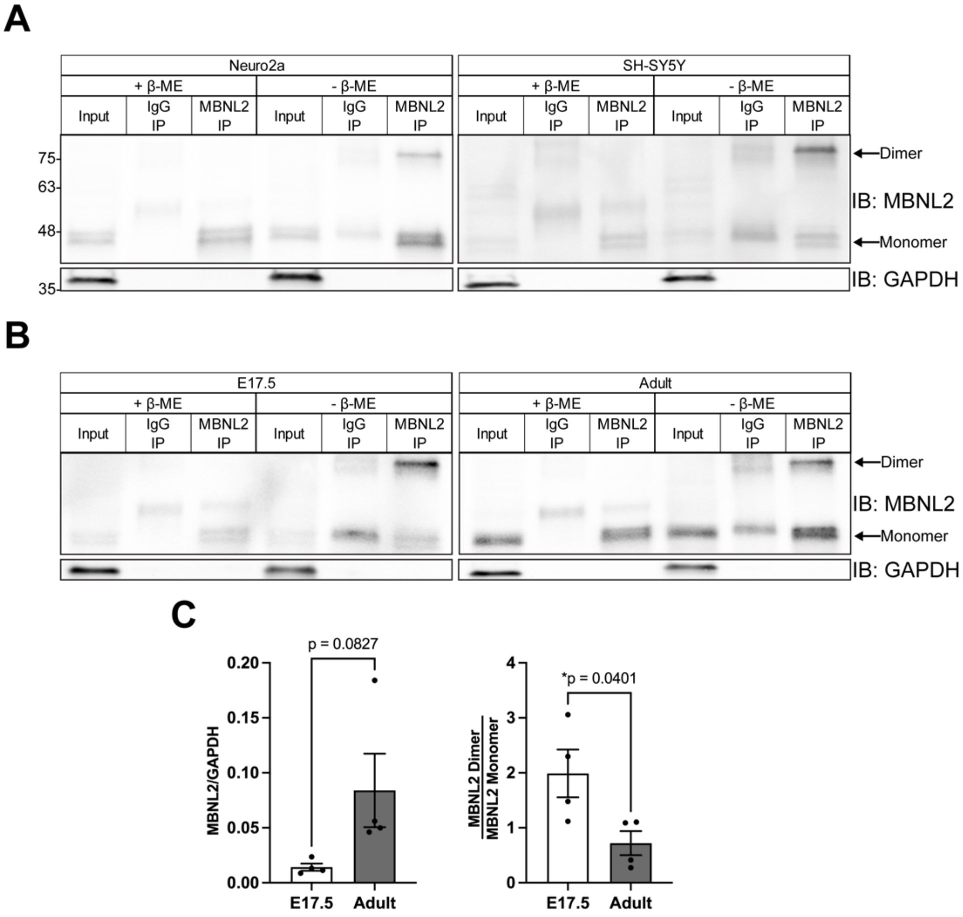


**Supplementary Figure 3.** MBNL2 forms dimers in mouse and human cell lines and embryonic and adult mouse brain tisssue. (**A**) Representative western blots of endogenous MBNL2 immunoprecipitated from Neuro2a and SH-SY5Y cells. Samples were prepared or eluted using Laemmli buffer containing or lacking β-ME. (**B**) Representative western blots of endogenous MBNL2 immunoprecipitated from E17.5 and adult mouse whole brain lysate. Samples were prepared or eluted with or without β-ME. (**C**) Quantification of MBNL2 levels normalized to GAPDH (left) and the dimer-to-monomer ratio of immunoprecipitated MBNL2 eluted without β-ME (right). Mean and standard error are reported across 4 biological replicates. *$p<0.05$ by two-tailed unpaired t test.

#### Supplementary Figure 4

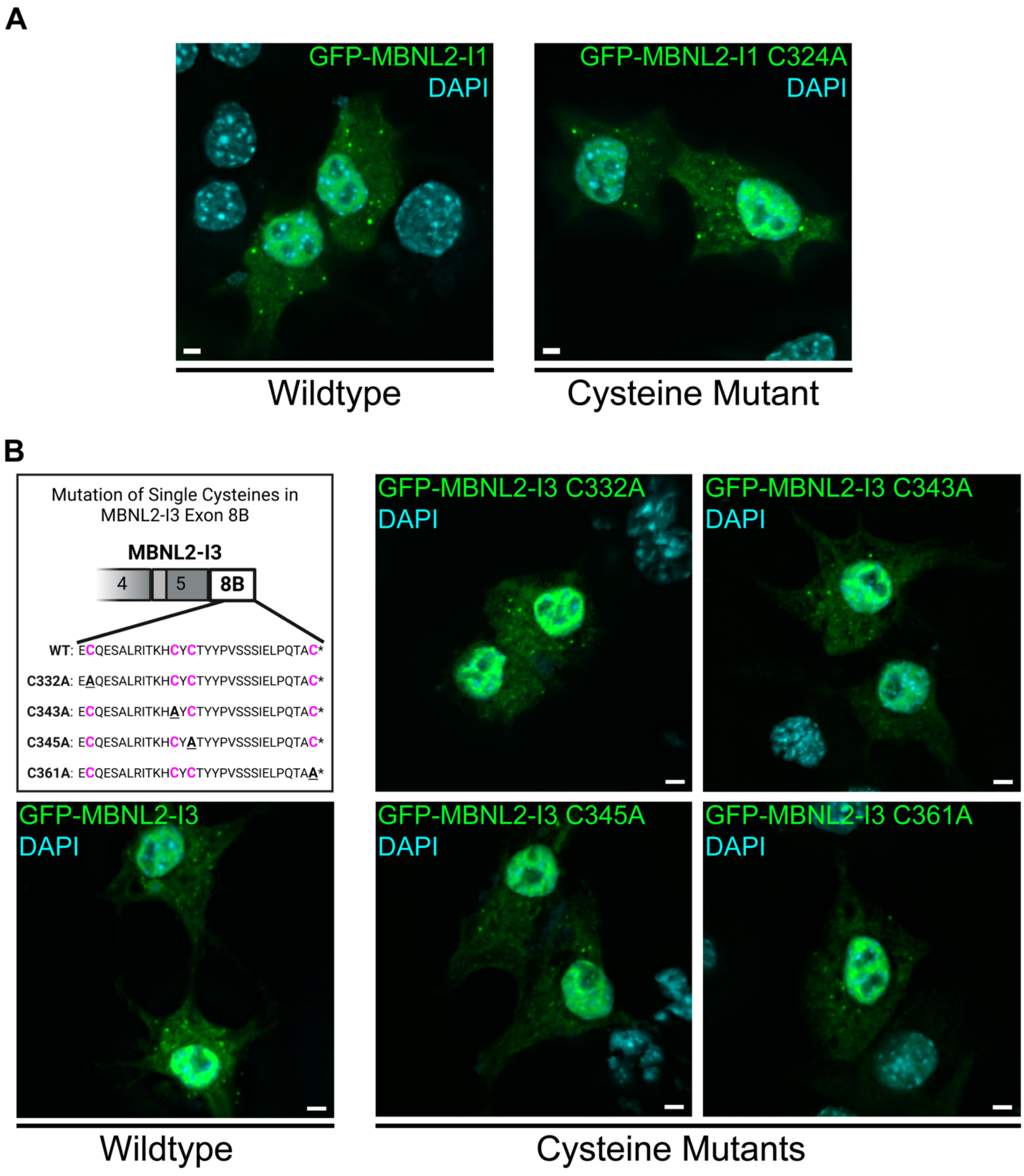


**Supplementary Figure 4.** Mutation of cysteines located in C-terminal domain of MBNL2 does not alter its distribution. (**A**) Representative images depicting the distribution wildtype and C324A versions of GFP-MBNL2-I1 when overexpressed in Neuro2a cells. Scale bars = 3µm. (**B**) (Top left) Illustration of the MBNL2-I3 C-terminal domain with the location of cysteines and C→A mutation in each MBNL1-I3 cysteine mutant. Created in BioRender. Bassell Lab, G. (2025) https://BioRender.com/y89o916. (Bottom left) Representative image of Neuro2a cells transfected with wildtype GFP-MBNL2-I3 and (Right) its associated cysteine mutants (C332A, C343A, C345A, and C361A). Scale bars = 3μm

#### Supplementary Figure 5

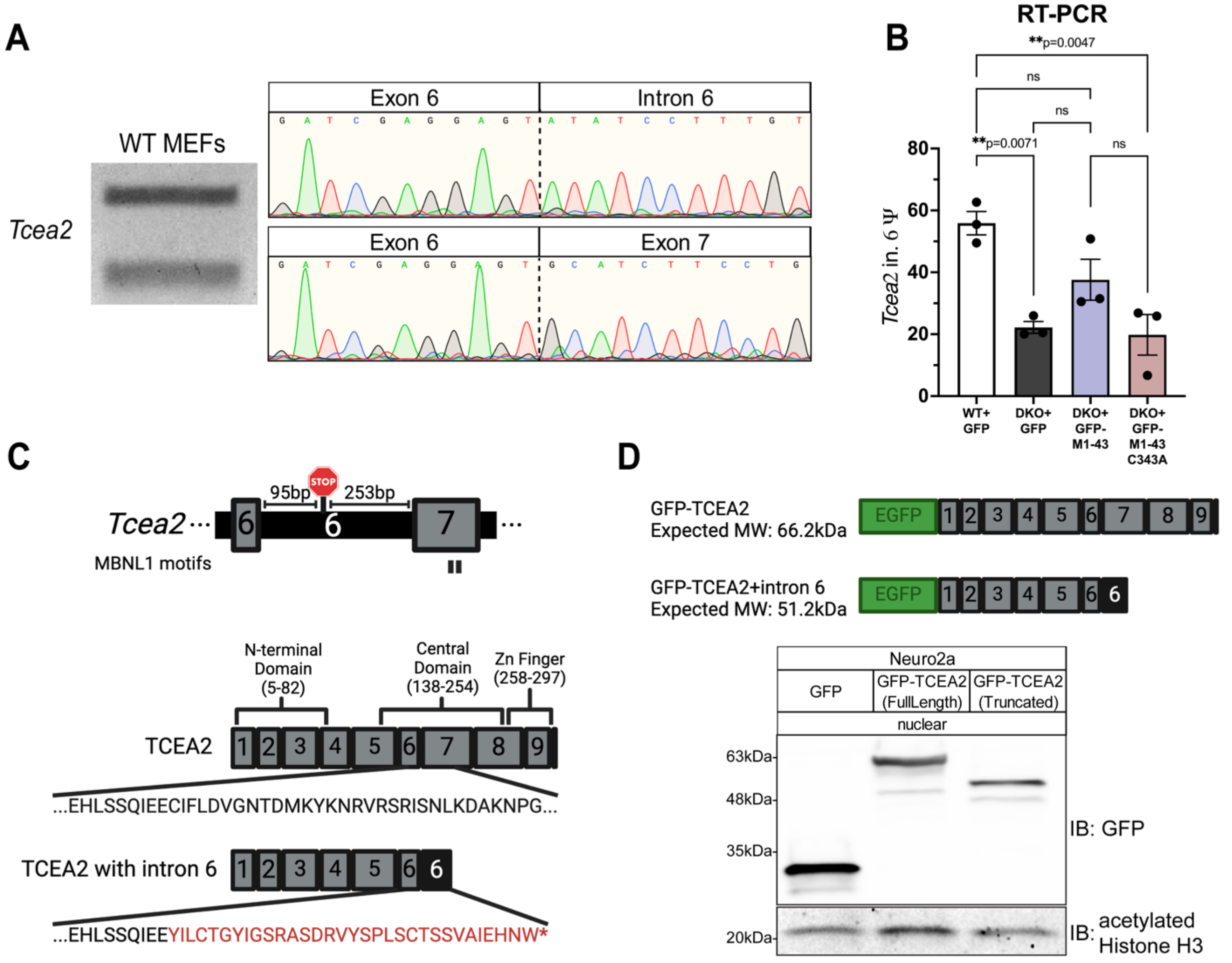


**Supplementary Figure 5.** Sanger sequencing confirms retention of *Tcea2* intron 6, which is also sensitive to dimerization of nuclear isoform MBNL1-43. (**A**) Sanger sequencing chromatograms of the *Tcea2* splicing products generated by RT-PCR on RNA from WT MEFs. The location of the Exon 6-Intron 6 and Exon 6-Exon 7 junctions for the upper and lower band, respectively, are indicated by dotted lines. (**B**) Quantitation of *Tcea2* intron 6 Ψ values from RT-PCR splicing gels using RNA from WT MEFs, DKO MEFs, and DKO MEFs transfected with either WT or C343A GFP-MBNL1-43. Data is reported as mean and standard error of 3 biological replicates. $^{**}p<0.01$ by one-way ANOVA with Tukey's post-hoc test. (**C**) (Above) Schematic depicting the location of a premature stop codon in intron 6 of *Tcea2*. (Below) Illustration of the structure of canonical TCEA2 protein along with the location of important functional domains and the neopeptide tail (red) produced by the retention of intron 6. Created in BioRender. Bassell Lab, G. (2025) https://BioRender.com/i05v148. (**D**) (Above) Schematic of the full-length and truncated versions of TCEA2 cloned into a N-terminal GFP vector, along with their expected molecular weights. Created in BioRender. Bassell Lab, G. (2025) https://BioRender.com/i05v148. (Below) Representative western blot of GFP and the full-length and truncated versions of GFP-TCEA2 from the nuclear fraction of transfected Neuro2a cells.

#### Supplementary Figure 6

A

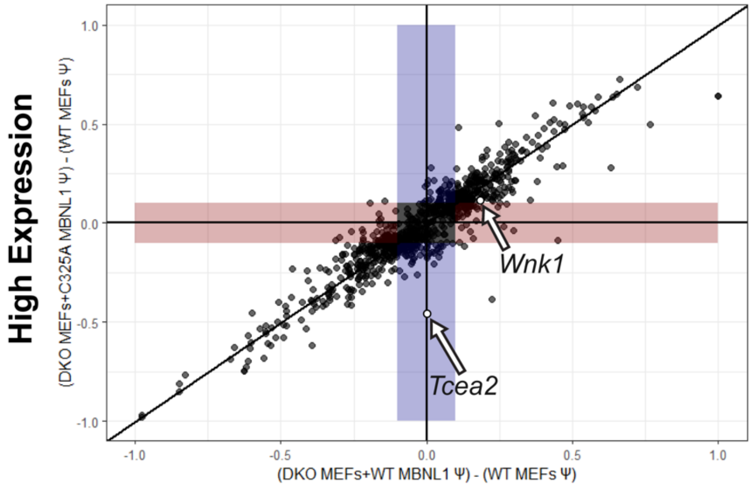


B

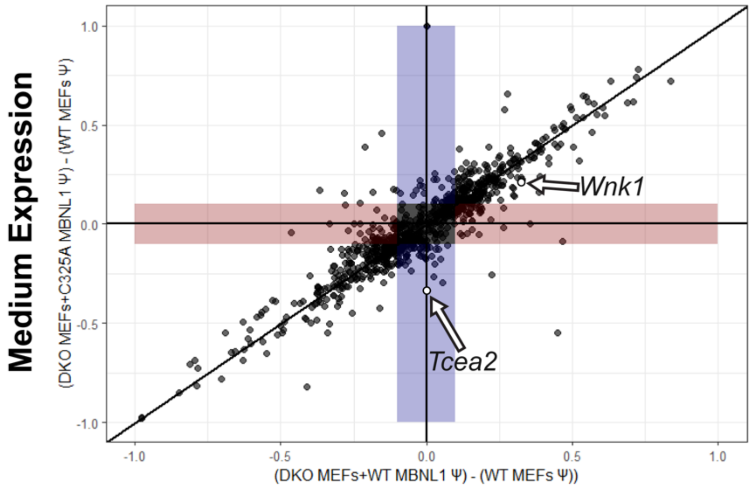


C

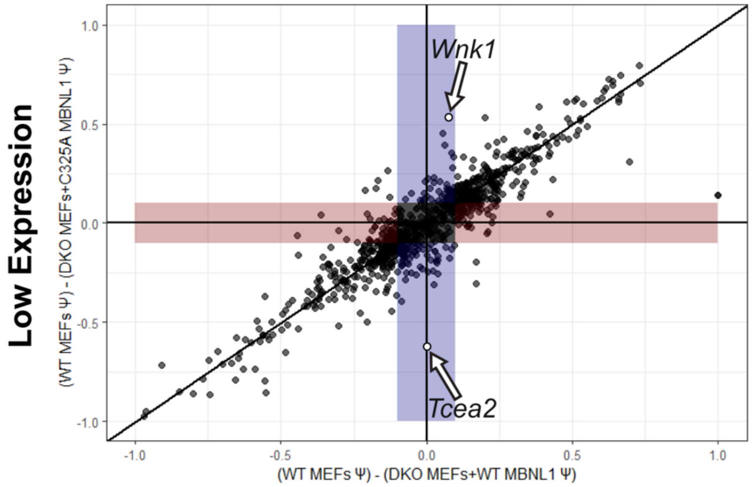


D

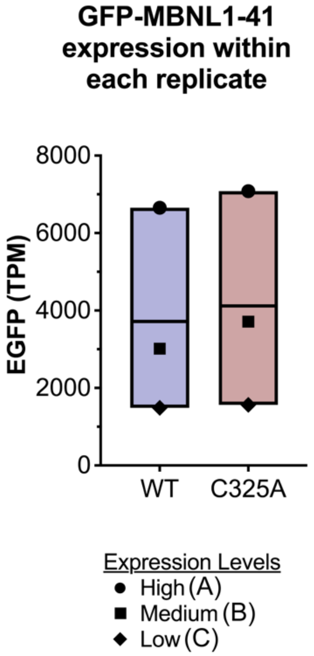


E

**Splicing rescue efficiency of GFP-MBNL1-41 expression levels within each replicate**

| EGFP Expression Level | *Tcea2* intron 6 | |
|---|---|---|
| | $\Psi_{DKO\ MEF+GFP\text{-}MBNL1\text{-}41}-\Psi_{WT\ MEF}$ | |
| | WT | C325A |
| Average | **0** | -0.453 |
| High | **0** | -0.400 |
| Medium | **0** | -0.333 |
| Low | **0** | -0.625 |

**Rescue**: $|\Psi_{DKO\ MEF+GFP\text{-}MBNL1\text{-}41}-\Psi_{WT\ MEF}| \leq 0.1$

| EGFP Expression Level | *Wnk1* ex.12 | |
|---|---|---|
| | $\Psi_{DKO\ MEF+GFP\text{-}MBNL1\text{-}41}-\Psi_{WT\ MEF}$ | |
| | WT | C325A |
| Average | 0.184 | 0.289 |
| High | 0.153 | 0.102 |
| Medium | 0.325 | 0.211 |
| Low | **0.074** | 0.553 |

**Rescue**: $|\Psi_{DKO\ MEF+GFP\text{-}MBNL1\text{-}41}-\Psi_{WT\ MEF}| \leq 0.1$

**Supplementary Figure 6.** Dot plots comparing splicing rescue efficiency in DKO MEFs expressing WT versus C325A GFP-MBNL1-41 at different levels in each RNAseq replicate**.**
(**A**) Dot plot comparison of ΔΨ from WT MEF levels for MBNL-dependent ASEs when WT versus C325A GFP-MBNL1-41 is expressed in DKO MEFs at comparatively "high"; (**B**) "medium"; and (**C**) "low" levels, (**D**) as determined by EGFP TPM in each RNAseq replicate. Retention of *Tcea2* intron 6, which is an MBNL1 dimerization-dependent ASE (see **Figure 3**), is labeled. Skipping of *Wnk1* exon 12, which is MBNL1-dimerization and -dosage dependent (see **Figure 4C,D**), being rescued only by WT GFP-MBNL1-41 in the "low" level replicate, is also labeled. (**E**) Tables listing the ΔΨ values of *Tcea* 2 intron 6 (above) and *Wnk1* exon 12 (below) for different levels of WT versus C325A GFP-MBNL1-41 in each replicate, as well as the average of all three replicates (see **Figure 4A**). These values correspond to the coordinates of the labeled dots in each respective dot plot. Bold values indicate a rescue of the ASE to WT MEF levels (|ΔΨ|≤0.1).

### Supplementary Figure 7

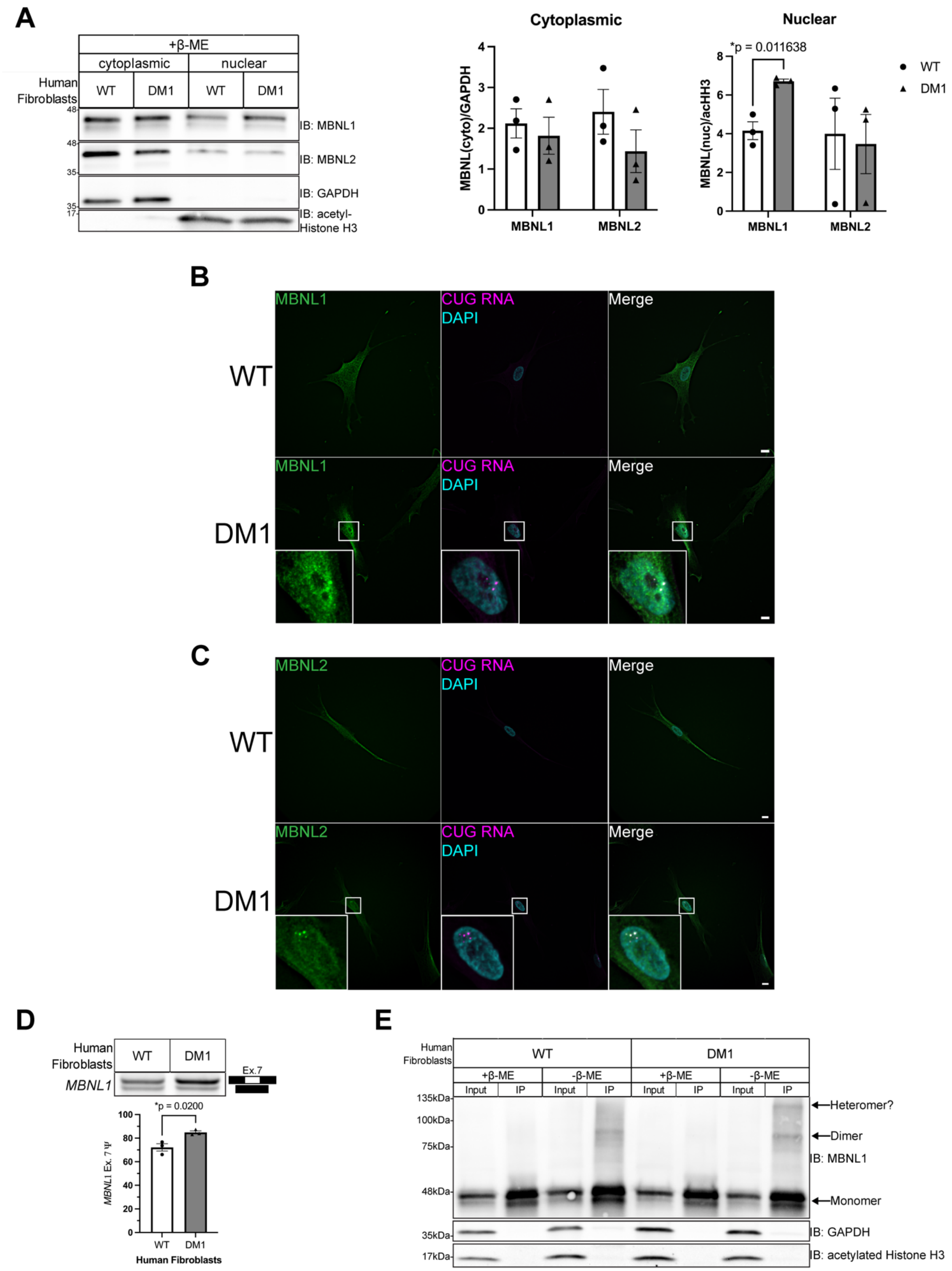


**Supplementary Figure 7.** MBNL1 forms higher order species in DM1 patient fibroblasts.

(**A**) (Left) Representative western blot against MBNL1, MBNL2, GAPDH, and acetylated Histone H3 nucleocytoplasmic fractionation of WT and DM1 patient fibroblasts. (Right) Quantification of cytoplasmic MBNL1/2 levels normalized to GAPDH (center) and nuclear MBNL1/2 levels normalized to acetyl-Histone H3 (acHH3; far right) in WT and DM1 patient fibroblasts. Mean and standard error are reported from 3 biological replicates. $^*p<0.05$ by two-tailed unpaired t test. (**B**) Representative FISH/IF images for CUG repeat RNA (magenta) and either MBNL1 (green) or (**C**) MBNL2 (green) protein in WT and DM1 patient fibroblasts. Nuclei are labeled with DAPI (blue). Scale bars = 10μm (**D**) (Above) Representative RT-PCR splicing gel for exon 7 of *MBNL1* conducted on RNA from WT and DM1 patient fibroblasts. (Below) Quantitation of *MBNL1* exon 7 Ψ values from RT-PCR splicing gels. Data is reported as mean and standard error of 3 biological replicates. $^*p<0.05$ by two-tailed unpaired t test. (**E**) Representative western blot of endogenous MBNL1 immunoprecipitated from WT or DM1 patient fibroblasts. Samples were prepared or eluted in Laemmli buffer with or without β-ME.

#### Supplementary Figure 8

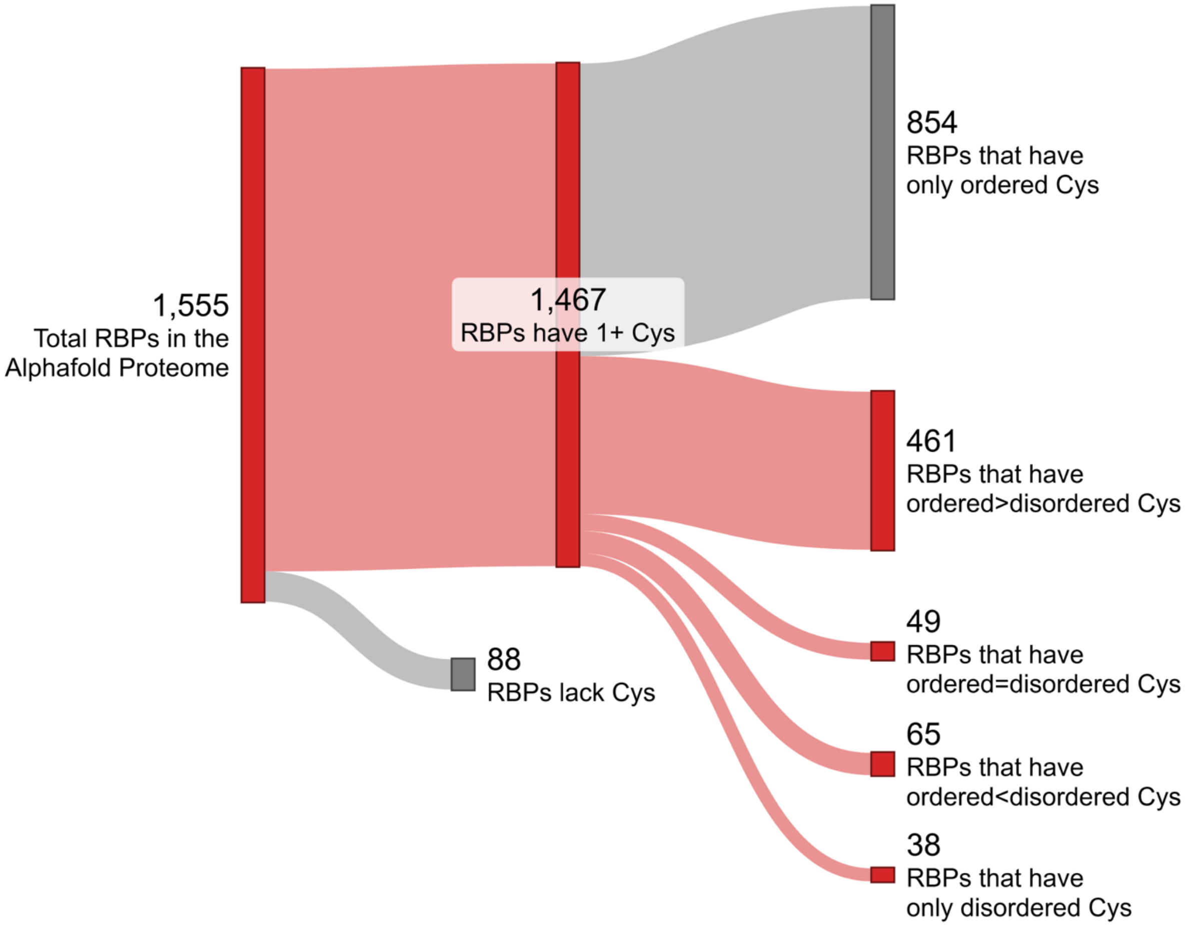


**Supplementary Figure 8.** Sankey plot depicting the proportion of RBPs that contain cysteine residues and the proportion of those RBPs whose cysteine residues are located in ordered or disordered regions of the protein. For reference, the initial list of RBPs analyzed was taken from (1) and narrowed down to Ensembl canonical transcripts. Classification of cysteine residues as located in either ordered or disordered regions is based on a per-residue confidence metric, called pLDDT, produced by protein structure prediction program AlphaFold (2,3). A pLDDT value less than 50 denotes a residue that is disordered in nature (4). The Sankey plot was created using SankeyMATIC (sankeymatic.com).
